# Multi-omics characterization of breast cancer metabolism identifies new metabolic targets

**DOI:** 10.64898/2026.03.06.710028

**Authors:** Hanneke Leegwater, Xiaobing Zhang, Luojiao Huang, Christine Hoegen, Xuesong Wang, Agnieszka B. Wegrzyn, Thomas Hankemeier, Bob van de Water, Erik Danen, Annelien J. M. Zweemer, Amy C. Harms, Sylvia E. Le Dévédec

## Abstract

Breast cancer cells undergo metabolic reprogramming to support proliferation and metastasis, but these alterations are heterogeneous. To systematically assess this heterogeneity under controlled conditions, we profiled 31 intracellular polar metabolites and 50 amines, together with exchange rates of 57 amines, in a panel of 51 breast cancer cell lines spanning diverse phenotypes. These data were integrated with lipidomics and transcriptomics. Multi-omics factor analysis identified metabolic signatures linked to proliferation and to differences between luminal, HER2-positive, basal A, and basal B cell lines, overlapping with an epithelial-to-mesenchymal transition phenotype. Fast-proliferating, aggressive cell lines showed increased uptake of essential amino acids and altered nucleotide levels, while heterogeneity in glutamine metabolism was mainly driven the subtype. This was partially associated with heterogeneous expression of metabolite transporters. To test the functional relevance, 67 metabolic genes were silenced in Hs578T using siRNA, and effects on proliferation and migration were measured by a sulforhodamine B assay, live-cell imaging, and a random cell migration assay. This resulted in 34 knockdowns that reduced proliferation and 20 reduced migration, with strong effects for the glycosyltransferases EXT1 and EXT2, nucleotide metabolism genes GART and HPRT1, and metabolite transporters SLC7A1, SLC7A11, and SLC16A3. These findings highlight phenotype-specific metabolic dependencies and identify candidate drug targets in aggressive breast cancer.

## Introduction

Metabolism is altered in cancers to sustain rapid proliferation. These alterations include enhanced energy production and use, increased nucleotide synthesis, and an increased dependence on amino acids, for protein synthesis or for *de novo* metabolite synthesis (Chandel et al., 2025; Hanahan & Weinberg, 2011; Pavlova & Thompson, 2016). Altered metabolism also contributes to therapy resistance (Zaal & Berkers, 2018) and metastasis (Bergers & Fendt, 2021; Elia et al., 2018). This increased dependency on specific metabolic processes has been proposed as a therapeutic strategy, to weaken cancer cells and improve combination therapies.

Metabolic alterations in breast cancer are heterogeneous and depend on the subtype and tumor stage (Demicco et al., 2024; Tan & Le, 2021). Some metabolic changes are driven by hormone receptors, such as the estrogen receptor (ER), progesterone receptor (PR), and HER2 receptor, but more specific mutations can lead to alterations in metabolism as well (Tan & Le, 2021). As a result, breast cancers can be distinguished from healthy tissue, and hormone receptor positive and negative subtypes can be recognized based on metabolic characteristics (Porcari et al., 2018).

Amino acid metabolism is an example of breast cancer-associated metabolic reprogramming. This includes changes in serine biosynthesis, due to altered phosphoglycerate dehydrogenase (PHGDH) expression, which can drive breast cancer metastasis (Rossi et al., 2022; Tan & Le, 2021). Glutamine metabolism is also rewired. Glutamine serves as a substrate for protein, hexosamine, and nucleotide synthesis, and as an anaplerotic substrate to the tricarboxylic acid (TCA) cycle (Pavlova et al., 2022). Elevated glutamine metabolism can be driven by the oncogene Myc (Wise et al., 2008). Triple negative breast cancer (TNBC) cell lines appear to depend more on glutamine metabolism than luminal A cell lines (Wahi et al., 2024), and patients with TNBC have elevated glutaminolysis compared to hormone-positive breast cancer patients (Cao et al., 2014). However, other studies report that HER2-positive cancers (Kim et al., 2013), or advanced cancers independent of receptor status, such as endocrine-resistant ER-positive tumors (Demas et al., 2019), show the strongest glutamine dependence. These findings highlight the central role of amino acid and glutamine metabolism in breast cancer, while also emphasizing metabolic heterogeneity between subtypes.

Cell lines are widely used model systems to study cancer characteristics *in vitro*. Cancer cell lines were originally isolated from breast cancer patients and have been maintained in laboratories (Dai et al., 2017), but still mimic properties of tumors, including differences in hormone receptor expression. Cancer cell lines vary in proliferation rate (Yizhak et al., 2014) and metastatic potential (Jin et al., 2020). Applications of cell lines include examining the efficacy of drugs or gaining fundamental insights in tumor biology. Importantly, they can be used to study cancer heterogeneity with differences in therapy response between cell lines, even when these share receptor targets (Holliday & Speirs, 2011), and can therefore serve as model systems to study metabolic heterogeneity.

We and others have previously shown that breast cancer cell lines have different metabolic gene expression signatures associated with an epithelial-mesenchymal state and hormone receptor status (Cherkaoui et al., 2022; Koedoot et al., 2021; Leegwater et al., 2025; Shaul et al., 2014; Wahi et al., 2024). However, gene expression alone cannot fully explain all metabolic characteristics, necessitating multi-omics approaches that include direct metabolite measurements. Omics data integration often requires using data from multiple sources or laboratories (Cherkaoui et al., 2022; Li et al., 2019). A limitation with this is that differences in culture conditions could lead to artificial differences between datasets, since culturing cell lines in different media influences intracellular metabolism (Abbott et al., 2023; Ackermann & Tardito, 2019; Katzir et al., 2019). In addition, inconsistencies in the behavior of the same cell line between laboratories have been reported as well (Dai et al., 2017; Kleensang et al., 2016; Osborne et al., 1987). Minimizing laboratory- and medium-based variability is therefore essential in multi-omics studies.

In this study, we investigated the relationship between metabolism and intrinsic breast cancer phenotypes. We collected multiple metabolomics measurements on a panel of breast cancer cell lines with a wide range of phenotypic characteristics, including differences in proliferation rate, hormone receptor status, migration capacity, and tumor origin, thereby modeling a wide range of patient phenotypes *in vitro*. All cell lines were cultured in the same media, and all metabolomic data was generated in parallel with our previously described lipidome dataset (Leegwater et al., 2025). For data integration, we used transcriptome (Koedoot et al., 2021) data of the same cell line panel cultured under identical conditions in our laboratory, in order to minimize between-lab variability. Based on this multi-omics integration, we identified potential metabolic vulnerabilities, and used siRNA knockdowns to assess the effect of a set of candidate genes on cell proliferation and migration, leading to the identification of potential novel metabolic drug targets.

## Methods

### Cell culture

We selected 52 breast cancer cell lines with differences in phenotypic characteristics, including hormone receptor status, morphology, and proliferation rate, as described previously (Leegwater et al., 2024, 2025). The cell lines were obtained from the Erasmus MC, the Netherlands, and stored in liquid nitrogen. To minimize external metabolic influences, all cell lines were cultured in RPMI 1640 medium (containing 2 mM L-glutamine, 10 mM D-glucose, and no sodium pyruvate) supplemented with 10% fetal bovine serum (FBS), 25 IU/mL penicillin, and 25 µg/mL streptomycin at 37 °C, with 5% CO_2_. Cells were seeded in two parallel 10 cm dishes, each with 10 mL medium, using a starting density for each cell line that resulted in approximately 80% confluency at the time of collection. Media was refreshed 24 h after seeding and cells were collected after 72 h. Biological replicates were obtained by thawing a new vial of the same passage, cultured on a different day, for each cell line. One dish was used for cell counting, a bicinchoninic acid (BCA) assay, and previously reported lipidomics measurements (Leegwater et al., 2024, 2025). The second dish was used for metabolomics sample collection in this study.

### Sample collection and liquid chromatography-mass spectrometry

The sample collection protocols differed for adherent, suspension, and mixed adherent-suspension cell lines. For adherent cells, medium was collected and 1 mL medium was centrifuged at 233 rcf (1000 rpm) for 3 min at 4 °C to remove residual cells. From this, 500 µL supernatant was transferred to a new tube, snap-frozen in liquid nitrogen, and stored at −80 °C. The cells were washed three times with ice-cold saline solution (0.9 %, w/v) to remove remaining culture medium. 500 µL of dry ice-cold methanol/water solution (8:2, v/v) was added to each dish to quench the metabolism. Cells were scraped from each dish, snap-frozen in liquid nitrogen, and stored at −80 °C.

For suspension cells, the cell suspension, including both cells and culture medium, was centrifuged at 233 rcf (1000 rpm) for 3 min at 4 °C. 500 µL supernatant was collected, snap-frozen and stored at −80 °C. The cell pellets were washed three times with 5 mL ice-cold saline solution. Cell pellets were lysed in 500 µL dry ice-cold methanol/water solution (8:2, v/v). For mixed adherent-suspension cells, the adherent cell population was scraped in 500 µL methanol/water solution, and the suspension population was washed as described above. The suspension pellets were combined with scraped adherent cell lysates, to generate a single sample representing both the suspension and adherent populations.

#### Polar metabolomics – intracellular

To profile polar metabolites, 100 µL lysate was analyzed in two batches with a randomized injection order. Each batch design included blanks and pooled quality control (QC) samples, prepared by mixing 40 µL of each sample. These QC samples were analyzed every 10 injections to monitor data quality and to correct for instrument variability. The 100μL samples were spiked with an internal standard solution, containing isotope-labeled metabolites (1-^13^C, ^15^N-Isoleucine, U-^13^C_4_, U-^15^N_2_-Asparagine, 2,3,3-D_3_-Leucine, U-^13^C_4_, U-D_3_, 9-^15^N-Aspartate, U-^13^C_5_, U-D_5_, ^15^N-Glutamate, U-^13^C_5_-Alanine, 2,2-D_2_-Glycine, U-^13^C_5_-Glutamine, U-^13^C_5_-Valine, U-^13^C_6_-Lysine, U-^13^C_11_, U-^15^N_2_-Tryptophan, 2,2,3,3-D_4_-Succinate, 2,2,4,4-D_4_-Citric acid, ^15^N_5_-GMP, and U-^13^C_3_-Pyruvate). A double liquid-liquid extraction was performed with chloroform/methanol/water (1:1:1, v/v/v). The upper aqueous phase was collected and dried using a speedvac. The residue was reconstituted in methanol/water (1:1, v/v). Samples were transferred to vials and placed in an autosampler tray and cooled to 4°C until injection. 4 µL was injected into an HILIC-UPLC-triple time-of-flight (TripleTOF) system.

Chromatographic separation was performed on a SHIMAZU LC-30AD system with a SeQuant® ZIC-cHILIC HPLC Analytical PEEK column (Merck) with a flow of 0.2 mL/min using a 45 min gradient. The UPLC was coupled to electrospray ionization on a TripleTOF mass spectrometer (AB SCIEX TripleTOF 5600). Analytes were detected in the negative ion mode using TOF MS at high mass resolution.

#### Amine metabolomics – intracellular and metabolite exchange

To measure amines, 5 µL cell lysate or media sample was processed using the same batch design and QC strategy as for the polar metabolites. Samples were derivatized with AccQ-Tag as described previously (Noga et al., 2011). Briefly, samples were spiked with an internal standard solution containing ^13^C^15^N-labeled amino acids. Proteins were precipitated by adding methanol and the supernatant was dried using a speedvac. The residue was reconstituted in borate buffer (pH 8.8) with an AccQ-Tag AQC derivatization reagent (Waters). The samples were cooled to 4 °C until injection. 1 µL of the reaction mixture was injected into a UPLC-MS/MS system.

Chromatographic separation was performed using an Agilent 1290 Infinity II LC system with an AccQ-Tag Ultra column (Waters) with a flow of 0.7 mL/min over an 11 min gradient. The UPLC was coupled to electrospray ionization on a triple quadrupole mass spectrometer (AB SCIEX QTRAP 6500). Analytes were detected in the positive ion mode and monitored in Multiple Reaction Monitoring (MRM) using nominal mass resolution.

### Data processing and quality control

Data were processed using MultiQuant Software for Quantitative Analysis (AB SCIEX, v3.0.2). MRM or extracted MS peaks were integrated and normalized using internal standards as follows. If applicable, an ^13^C^15^N-labeled analog was used. For other metabolites, the closest eluting internal standard was used. mzQuality (Van Der Peet et al., 2025) was applied for batch correction and to correct for instrumental drift within a batch, based on the pooled QC samples. Amines and polar metabolites were included for further processing if the relative standard deviation of the quality control samples (RSDqc) was lower than 30%.

### Omics data preprocessing and normalization

#### Intracellular polar and amine metabolomics

Intracellular polar and amine metabolomic datasets were preprocessed using R v4.4.1 (R Core Team, 2024). Metabolites were reported as relative concentrations, which are peak areas divided by the peak area of an internal standard, to correct for deviations after metabolite extraction (amine target area/ISTD area; unit free). The two datasets were normalized separately to correct for the amount of measured sample using probabilistic quotient normalization (PQN) (Dieterle et al., 2006) and data was log_2_ transformed. Biological replicates were averaged, and the data was scaled for each metabolite to have mean 0 and standard deviation (SD) 1 across all cell lines. For metabolites detected on both platforms, the amine measurement was used in further processing steps.

#### Metabolite exchange rates

Absolute metabolite concentrations in media were estimated using peak areas per metabolite in each sample and a calibration line of seven defined concentrations per metabolite as described previously (Preciat et al., 2021; Van Der Peet et al., 2025). Metabolite exchange rates were calculated by subtracting the metabolite concentration in media after 48 h (C_48h_) by the mean metabolite concentration in twelve fresh media samples (C_0h_). The total dry weight per sample (DW) was estimated from a protein concentration measure of a parallel plate, determined previously by a BCA assay (Leegwater et al., 2024), and an estimated protein concentration of 0.7 g protein per gram dry weight (gDW) for all cell lines. These values were corrected for different growth rates by estimating the number of cells after 24 h, using the starting and final cell count numbers, assuming linear growth (DW_corrected_). To obtain metabolite exchange rates per cell line biological replicate in µL/gDW/h, the metabolite change was divided by 48 h and by the gram dry weight per cell line (formula 1). As a quality control step, cell line replicates were removed if all values for essential amino acids were secretions, which hints towards these cells having leaked their content into the media, or ‘negative’ cell growth, e.g. cell death. Based on this quality control step, all biological replicates for HCC1569 were removed. The metabolite exchange rate per cell line was the mean rate +/- SD of three biological replicates. For multi-omics integration in MOFA, exchange rates were scaled per metabolite to have a mean 0 and SD 1. For relative comparisons between metabolites, the rates were min-max normalized per metabolite, where 0 means no secretion or uptake, positive values are secretions, and negative values are uptakes, and the largest uptake or secretion was set to 1.

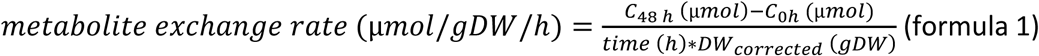

#### Lipidomics

Lipidomic data was obtained from a previous study, where data was normalized using PQN based on structural lipids, log_2_ transformed, autoscaled and averaged per lipid class (Leegwater et al., 2024, 2025).

#### Pathway score calculations

Pathway scores were calculated based on log_2_ transformed autoscaled gene expression values, obtained from a previous study in our group (Koedoot et al., 2021). First, genes with zero expression across all cell lines were removed. Gene identifiers for each metabolic pathway were obtained from Reactome v91 (https://zenodo.org/records/14518331, (Milacic et al., 2023). All sub-pathways under the top-pathway metabolism were included (R-HSA-1430728). The pathway score was calculated as the mean autoscaled gene expression per pathway, resulting in 273 pathway scores.

### Multi-omics data integration

Multi-omics factor analysis (MOFA) was performed to find shared variation in the omics datasets. MOFA is an unsupervised statistical approach to learn shared factors between omics datasets (Argelaguet et al., 2018). All input datasets, which were amine metabolites, polar metabolites, lipid class averages, and mRNA gene expression as pathway scores, were filtered to keep only cell lines measured in all datasets, which left 49 cell lines. Variance explained per factor, factor values per cell line and feature weights per factor were extracted from the MOFA object and further processed to find associations between cell line phenotypes and the factors. Correlations between MOFA factors and phenotypes were estimated using spearman correlations (r_s_).

### Correlations between MOFA factors and metabolite transporters

To define genes that can transport metabolites of interest, a generic human metabolic reconstruction, Recon3D v3.01, was downloaded from the Virtual Metabolic Human database (VMH) database (Brunk et al., 2018); https://vmh.life). The model was loaded in MATLAB R2022b, and by using the Cobra Toolbox (v3.0, (Heirendt et al., 2019), all genes were extracted if they belonged to a reaction from the subsystem ‘Transport, extracellular’. To filter transport reactions for metabolites with high weights in the MOFA, extracellular metabolite identifiers were obtained from the VMH website and reactions were selected if they imported or exported at least one of the metabolites of interest, and were in the subsystem ‘Transport, extracellular’. A second set of metabolite transporters was obtained from Reactome v91, where genes related to ‘transport of small molecules’ (R-HSA-382551) were included.

We aimed to identify transporters, either defined by Recon3D or by Reactome, for which the expression correlated with MOFA Factor 1, Factor 2, proliferation rate, or with EMT score, which we previously defined as a numeric representation of the difference in basal-luminal breast cancer subtypes (Leegwater et al., 2025). Spearman correlations (r_s_) were calculated between each transporter’s log_2_ transformed gene expression and the numerical phenotypic data. Pearson correlations (r) were calculated between genes.

### Differential metabolic gene expression analysis

RNA sequencing data (Koedoot et al., 2021) was filtered to keep genes with at least one count per million (CPM) in five or more cell lines and a mean count higher than 0.1 CPM. DESeq2 was used to obtain differentially expressed genes associated with Factor 1 or cell line subtype, with a FDR < 0.1. To select genes positively associated with proliferation, the coefficient of Factor 1 was filtered to be > 0, and a log_2_ fold change of > 1 was used to determine upregulated genes in basal B breast cancer cell lines compared to luminal cell lines. Basal A and HER2-positive cell lines were included in the model fit to improve dispersion estimates.

Data on Breast Invasive Carcinoma patients and breast cancer subtype classifications from The Cancer Genome Atlas project were obtained from cBioPortal (https://www.cbioportal.org/datasets, TCGA, PanCancer Atlas) on 08-04-2021 (Cerami et al., 2012; Hoadley et al., 2018). Between-sample-normalized RNA sequencing counts were filtered to keep only female patients with a defined breast cancer subtype, and all normal breast tissues samples. DESeq2 was used to identify genes elevated in basal breast cancer compared to normal breast tissue (log_2_ fold change > 0, FDR < 0.1).

#### Definition of metabolic genes

A gene was defined as “metabolic gene” if it was listed in at least two of the following resources: (1) genes in a reaction in the genome-scale metabolic model Recon3D v3.01 (Brunk et al., 2018), (2) genes annotated to the gene ontology term ‘metabolic process’ (GO:0008152, GO version 2024-01-17; https://zenodo.org/records/10536401 (Ashburner et al., 2000; The Gene Ontology Consortium et al., 2023)), (3) genes annotated to the human KEGG ‘Metabolic pathways’ (hsa01100, release 114.0+/05-19, may 2025; (Kanehisa et al., 2024)) or (4) genes in Reactome ‘metabolism’ or ‘transport of small molecules’ (R-HSA-382551 and R-HSA-1430728, v91). Genes involved in both metabolism and other biological processes were also included, for example in signaling or post-translational modification if a phosphate group is involved, or in protein complexes, such as the proteasome or ribosome.

#### Network analysis of metabolic targets

To visualize metabolic-subprocesses of differentially expressed genes, genes were mapped to the STRING network (v12.0 on 16-05-2025 (Szklarczyk et al., 2023)), with Homo sapiens as organism, based on the full STRING network, a medium confidence score of 0.400 and medium FDR stringency (5 percent). The resulting network was exported to Cytoscape (v.3.10.1). In Cytoscape, the network was clustered using MCL clustering with the Cytoscape plugins stringApp (v2.2.0; (Doncheva et al., 2019)), clusterMaker2 (v2.3.4 (Morris et al., 2011)) and yFiles Layout Algorithms (v1.1.5). Nodes were colored, based on Reactome pathways and STRING clusters, obtained from STRING enrichment within the stringApp.

### Data visualization

Scaled data and correlation matrices were visualized as heatmaps using ComplexHeatmap (Gu, 2022). If clustering was used, this was hierarchical clustering, using Euclidean distance and complete linkage. MOFA2 was used to visualize variance after multi-omics factor analysis. Scatter plots and lollipop plots generated with ggplot2 and cowplot in R (Wickham, 2016; Wilke, 2020). DrawIO (https://www.drawio.com/) was used to draw flow charts.

### Small interfering RNA knockdown screen

Transient siRNA-mediated knockdowns were performed in the cell line Hs578T as described previously (Koedoot et al., 2019). Briefly, transient siRNA knockdowns were performed in 96-well plates by reverse transfection of 50 nM SMARTpool siRNA (Dharmacon, Lafayette, CO, USA), which contains four different siRNAs, using the transfection reagent INTERFERin (Polyplus, Illkirch, France) according to the manufacturer’s guidelines. The following controls were included during transfection: transfection reagent only (Mock), a mixture of 720 siRNAs targeting different kinase genes (non-specific kinase pool, siKinasePool), and siRNA against integrin β, which should reduce cell migration. Medium was refreshed after 24 h and transfected cells were seeded after 48 h into separate plates for different phenotypic assays.

#### Random cell migration assay

Cells were seeded in collagen-coated plates and stained with Hoechst-33342 (100 ng/mL) after 24 h. Treatment with 1 µM flavopiridol or 1 µM bosutinib was included as positive control for migration inhibition. Imaging of Hoechst-33342 signals was performed every 15 minutes for 16 h using a 20X objective on an ImageXpress Micro XLS imager (Molecular Devices). These images were analyzed as described previously (Koedoot et al., 2019), resulting in a migration speed per cell line replicate. The speed was averaged for three biological replicates and reported as the mean speed in µm/h with the standard error of the mean (SEM).

#### Cell proliferation

To measure cell proliferation, a sulforhodamine B (SRB) assay, as described previously (van der Noord et al., 2023; Vichai & Kirtikara, 2006), and a confluency assay were performed 24h after reseeding. Treatment with 1 µM flavopiridol or 1 µM bosutinib was included as positive controls. Confluency was measured using an IncuCyte S3 microscope (Sartorius) and analyzed using IncuCyte 2020B with settings optimized to distinguish Hs578T confluency. Images were collected every 4 h. After 72 h, the plates were fixed using 50% trichloroacetic acid (TCA, 30 µL in 100 µL medium, Sigma-Aldrich) and stained with 0.4% SRB and compared to the 0 h timepoint. For each biological replicate, the IncuCyte percentage confluency was subtracted from the percentage confluency of the mock control. For the SRB data, data for each biological replicate was min-max normalized, where 0 represents the concentration at T0, and 100 the concentration of the mock after 72 h.

#### siRNA knockout screen statistical analysis

To calculate changes in migration and proliferation were significant, a two-sided t-test was performed with unequal variance for each siRNA compared to the mock control, and an FDR < 0.1 was used to correct for multiple testing. A change of 20% in migration and either 20% or 50% in proliferation was considered as a biologically relevant effect size.

### Data and code availability

Processed, normalized metabolomics can be found in **Supplementary Table 1**. Metabolomics raw data is available upon request. All code to reproduce these analyses are available on GitHub (https://github.com/LACDR-CDS/Leegwater_metabolomics) and archived to Zenodo (link will be added when the manuscript is accepted for publication).

## Results

### Metabolism links proliferation and breast cancer subtypes

To map the metabolic heterogeneity in breast cancer cell lines, metabolites were measured in 51 breast cancer cell lines. After data preprocessing, normalization, and quality control, 31 intracellular polar metabolites and 50 amines were detected and quantified, and metabolite uptake and secretion rates were determined for 57 amines. These data were integrated with previously published lipidomic data, which included 21 lipid classes (Leegwater et al., 2025), and with gene expression pathway activity scores for 273 metabolic pathways in 49 cell lines, calculated based on the Reactome database (Milacic et al., 2023) and previously published transcriptomic data (Koedoot et al., 2021).

Multi-omics factor analysis identified eight latent factors representing shared variation across the omics datasets. Factor 1 captured variation in all omics datasets, especially in metabolite uptake and secretion rates (**Figure 1A**). Factor 2 captured variation across all omics datasets except polar metabolomics, and other factors captured variation in one or a subset of omics datasets (**Figure 1A**).

**Figure 1.**
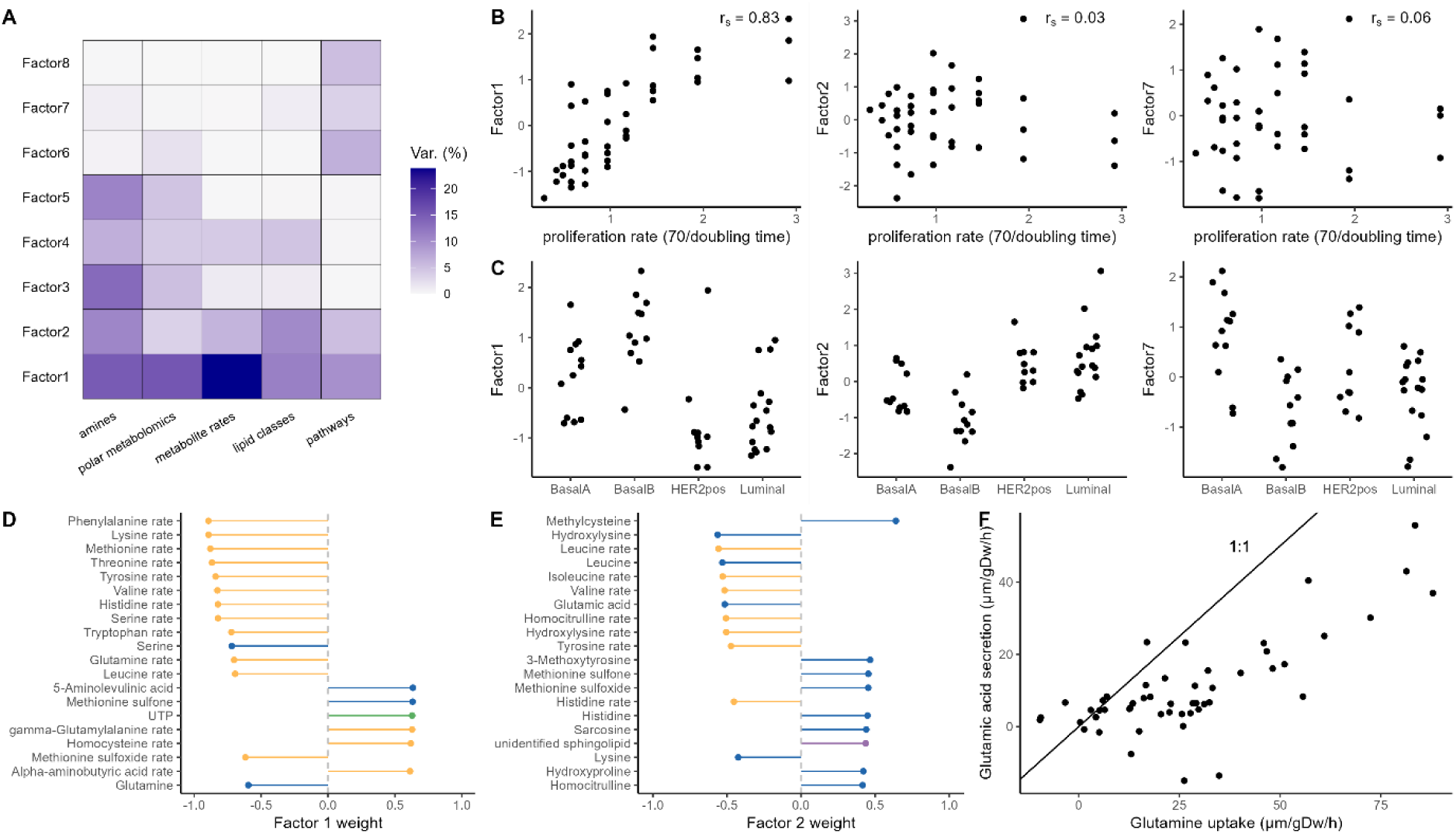
Variation in breast cancer cell line metabolism is related to proliferation and breast cancer subtypes. (A) Percent variance explained per omics dataset in each factor after multiomics factor analysis (MOFA) for a panel of 49 breast cancer cell lines. (B) Spearman correlation between proliferation rate for each cell line and MOFA factor scores 1, 2, and 7. (C) Relation between cell line MOFA factor scores 1, 2, and 7 and cell line subtypes. (D-E) Top scoring feature weight contributing to Factor 1 and 2. The features are colored by omics dataset, for amines (blue), polar mteabolomics (green), metabolite uptake and secretion rates (yellow), lipid classes (purple). A positive weight means a positive contribution to a factor. Because metabolite rates were scaled before omics integration, a positive contribution can mean a higher secretion, or a lower uptake rate, or a mix of both if some cell lines secrete a metabolite and others take it up. (F) Correlation between glutamine uptake and glutamine secretion in each cell line. The black line represents 1:1, which represents that one molar glutamine would be secreted as one molar glutamate.

To test whether any of the latent factors were associated with cell line phenotypes, we calculated if the factors were linked to proliferation rates and differences in cell line subtypes. Factor 1 correlated strongly with cell line proliferation rate (r_s_ = 0.83, **Figure 1B**). No other factors correlated with proliferation (|r_s_| < 0.3; **Figure 1B**). Since basal B cell lines, the most aggressive subtype in this breast cancer cell line panel, proliferate faster on average than other cell line subtypes, we examined whether Factor 1 also captured differences between subtypes. Indeed, HER2-positive and luminal cell lines were lower in Factor 1, whereas basal A and basal B cell lines were higher, overlapping with the fast-proliferative phenotype (**Figure 1C**). Factor 1, Factor 2, and Factor 7 captured significant differences between cell line subtypes basal A, basal B, HER2-positive, and luminal (ANOVA P < 0.001, **Figure 1C**), where Factor 2 separated basal B cell lines from other subtypes, and Factor 7 mainly separated basal A cell lines and a subset of HER2-positive cell lines.

#### Proliferation-associated metabolic variation

Features with high weights for the proliferation-associated Factor 1 included several metabolic processes. All essential amino acids (Lopez & Mohiuddin, 2025) showed increased uptake in fast-proliferating cell lines (phenylalanine, lysine, methionine, threonine, valine, histidine, tryptophan, leucine; **Figure 1D**, and isoleucine **Supplementary Figure 1**). As essential amino acids are required for protein synthesis but cannot be synthesized by cells, this indicates an increased use of environmental metabolites in fast-proliferating cell lines. Factor 1 also captured an increased uptake of tyrosine and serine, together with lower intracellular serine levels. Serine can be used for synthesis of proteins and the membrane lipid phosphatidylserine (PS). There was a small positive weight for the pathway “Synthesis of PS” and for intracellular PS levels (Factor 1 weights both 0.18), suggesting that serine contributes to both protein synthesis and PS synthesis in fast-proliferating cell lines.

Other metabolites contributed to Factor 1 as well. Increased secretion of γ-glutamylalanine and glutathione, higher intracellular glutathione, and increased uptake of methionine sulfone and methionine sulfoxide may reflect protein breakdown or oxidative stress. Succinate and succinyl-CoA were lower in fast-proliferating cell lines, but other TCA cycle intermediates did not differ, and neither did the gene expression pathways “Citric acid cycle (TCA cycle)”, “Maturation of TCA enzymes and regulation of TCA cycle”, and “Glycolysis”. This suggested that energy metabolism was not globally reprogrammed in fast-proliferating cell lines.

The fast-proliferating cell lines in Factor 1 had higher intracellular UTP, GTP, CTP, and ATP together with elevated pathways “pyrimidine biosynthesis” and “purine ribonucleoside monophosphate biosynthesis” (**Supplementary Figure 1** with top 50). UTP, GTP, ATP, and CTP are precursors for RNA and DNA, which could suggest increased rates of RNA and DNA synthesis in fast-proliferating cells, consistent with an increased demand for nucleotides.

#### Subtype-associated metabolic variation

Factor 2 highlighted differences specific to basal B cell lines. These included elevated hydroxylysine and leucine, along with altered leucine and hydroxylysine rates (**Figure 1D**). Multiple other metabolite rates were altered, including all three branched-chain amino acids, leucine, isoleucine, and valine, and the amine homocitrulline. This suggests that metabolite uptake is not only related to proliferation-associated metabolic processes, but also that different cell line subtypes have different metabolic needs. Factor 2 also included cholesterol esters, hexosylceramides, and an unidentified sphingolipid, in line with previously linked lipidomic differences linked to morphology (Leegwater et al., 2025).

Both Factor 1 and Factor 2 reflected heterogeneity in glutamine metabolism (**Figure 1D, 1E**). Fast-proliferating cell lines showed increased glutamine uptake, lower intracellular glutamine, higher intracellular glutamic acid, and increased glutamate secretion (Factor 1 weights −0.7, - 0.6, 0.3, and 0.4, respectively), but intracellular glutamic acid and glutamic acid secretion also contributed to Factor 2 (Factor 2 weights −0.51 and −0.31). This suggested that heterogeneity in glutamine metabolism was associated with both proliferation and the basal B phenotype. Enhanced glutamine uptake and glutamate release were closely related, although not all glutamine was secreted as glutamate (**Figure 1F**), consistent with prior findings on a cancer cell line panel with multiple cancer types (Jain et al., 2012). Pathway-level gene expression of ‘glutamate and glutamine metabolism’ was not altered in either factor (weights < 0.1), suggesting additional regulation beyond transcription.

Factor 7, separated basal A cell lines from basal B and luminal cell lines, with HER2-positive as intermediate (**Figure 1C**). This was due to multiple gene expression pathways but not due to metabolite levels, and captured biological processes within glycosaminoglycan metabolism and energy metabolism (**Supplementary Figure 1**). Together, these results show that proliferation and subtype differences are linked to distinct metabolic programs, motivating further study of gene expression.

### Aggressive breast cancer cell lines show increased essential amino acid uptake

Next, we studied the increased metabolite rates associated with Factor 1 in more detail to gain more insights into metabolic processes related to proliferation. Essential amino acids, as well as tyrosine, glutamine, serine, and methionine sulfoxide showed a gradient of increasing uptakes with increased proliferation rates and Factor 1 scores (**Figure 2A**), while homocysteine, γ-glutamylalanine, α-aminobutyric acid and glutathione were secreted (**Figure 2A**). Factor 1 had low weights for metabolic pathways related to essential amino acids, indicating that the increased amino acid uptake could not be attributed to pathway-wide differences in gene expression.

**Figure 2.**
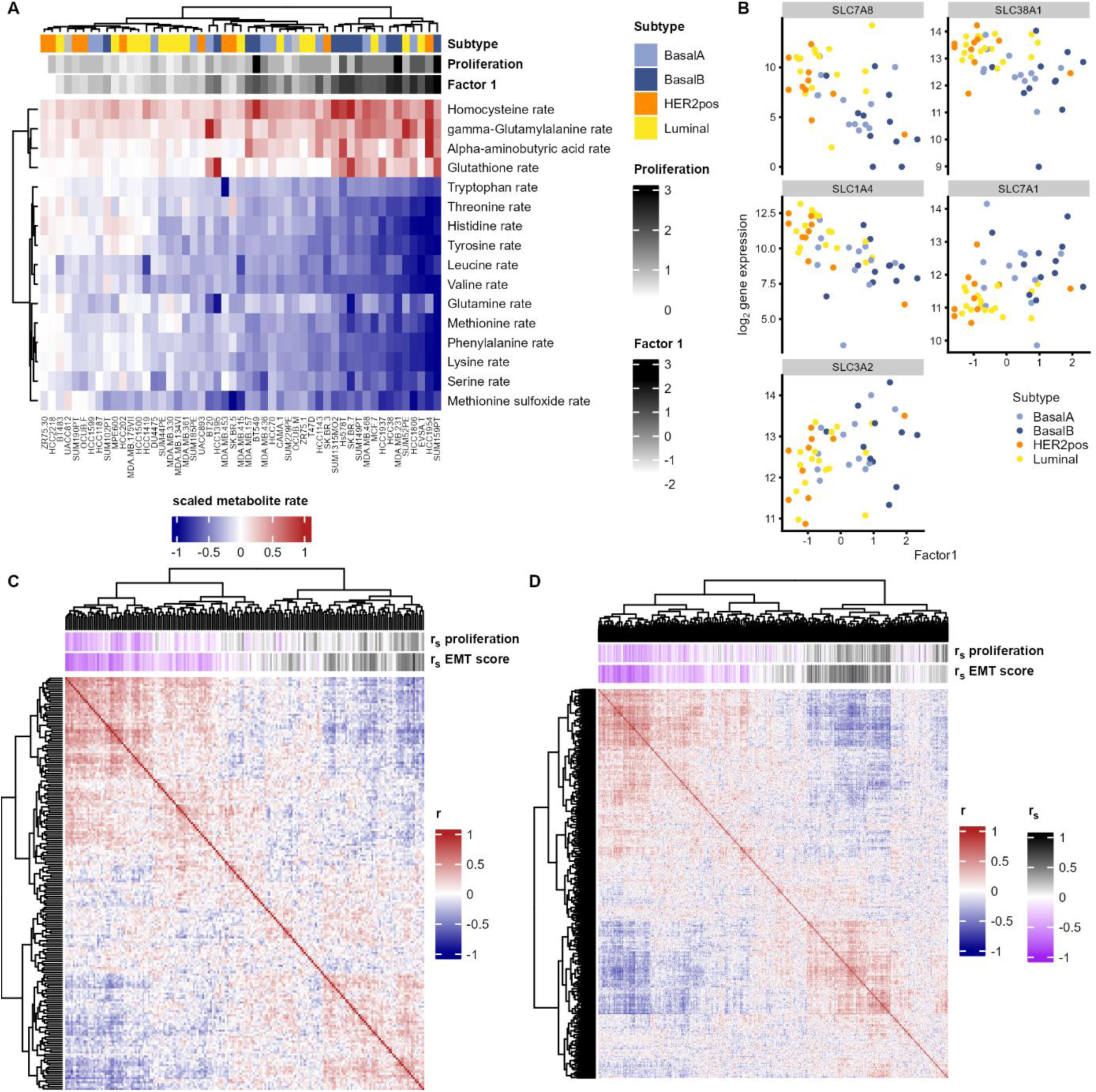
Breast cancer metabolite transport is linked to proliferation and subtype. (A) Metabolite uptake and secretion rates for metabolites associated with Factor 1 and proliferation. Metabolite rates are scaled with either −1 or 1 as maximum secretion or uptake, and 0 is zero uptake or secretion. (B) Gene expression for metabolite transporters that can transport one or more of the metabolites in A and are significantly associated with Factor 1. (C-D) Correlation heatmap of gene expression profiles of 191 and 565 metabolite transporters as defined by Recon3D (C) or by Reactome (D). The correlation heatmap is based on Pearson correlations. Metadata is colored based on Spearman correlations (r_s_) between gene expression profiles and proliferation rate or EMT score, where a higher EMT score means a more mesenchymal-like basal B phenotype.

Given this, we next considered other mechanisms that may facilitate increased uptake, such as enhanced activity of metabolite transporters. To test whether transporter activity contributed, gene expression profiles were analyzed for 35 transporters that transport the L-amino acid form of one or more of these amines. Of these, 24 transporters were expressed in this breast cancer panel (> 1 CPM in ≥ 5 cell lines) and 5 correlated with Factor 1 (|r_s_| > 0.4). Three neutral amino acid transporters (SLC1A4, SLC7A8, and SLC38A1) correlated negatively, meaning a lower expression in fast-proliferating cell lines, and two amino acid transporters (SLC3A2 and SLC7A1) correlated positively (**Figure 2B**).

To better understand the role of the five proliferation-associated metabolite transporters, gene expression profiles in cancer were compared to healthy tissue, based on data from The Cancer Genome Atlas (TCGA) (Hoadley et al., 2018; Liu et al., 2018). All five transporters had different gene expression profiles compared to healthy breast tissue (**Supplementary Figure 2**), but with a cancer subtype-specific effect: SLC1A4, SLC7A8, SLC38A1 were elevated in luminal and HER2-overexpressing cancers, SLC3A2 was elevated in all cancer subtypes, although less profound in luminal A breast cancers, and SLC7A1 was higher in basal cancers. This suggests that metabolite transporters are also mainly regulated by cell line molecular subtypes.

Next, we performed a broader gene expression analysis of metabolite transporters. Approximately two-thirds of the genes involved in “extracellular transport” (191 of 292; as defined by Recon3D) or “transport of small molecules” (565 of 869; as defined by Reactome,) were expressed in the cell line panel. Clustering by pairwise correlations revealed two major coexpressed transporter clusters (**Figure 2C**, **Figure 2D**). For each gene, the expression was correlated with the cell line’s matching proliferation rate or EMT score. We previously introduced the EMT score as a numeric interpretation of the difference between basal and luminal cell lines, since basal B cell lines in this panel have an overall mesenchymal-like phenotype and luminal cell lines have an epithelial-like phenotype (Leegwater et al., 2025; Rogkoti et al., n.d.). This showed that the two diverging clusters overlapped mainly with EMT score, but also with proliferation rate, meaning that fast-proliferating, mesenchymal-like cells had a different set of metabolite transporters expressed, which could contribute to the heterogeneity in metabolite uptake rates.

### Selecting metabolic targets in aggressive breast cancer cell lines

Having established that both proliferation and differences between basal B and luminal cell lines were linked to metabolite profiles and exchange rates, we next aimed to identify metabolic genes driving these aggressive phenotypes. Specifically, the aim was to select metabolic targets for functional studies to validate that our computational data mining could predict potential metabolic drug targets, focusing on targets whose modulation might reduce proliferation or reverse an aggressive migratory phenotype. Metabolic genes were selected if these were expressed in at least five cell lines (CPM > 1), and either elevated in basal B compared to luminal cell lines (log_2_ fold change > 1, FDR < 0.1) or positively associated with Factor 1 (coefficient > 0, FDR < 0.1; **Figure 3A-C**). This approach identified 806 metabolic genes. Since a higher expression in basal B compared to luminal cell lines does not distinguish basal B upregulation from luminal downregulation compared to healthy cells, we refined our selection by requiring that gene expression was elevated in basal breast cancer patients compared to healthy breast tissue (log_2_ fold change > 0, FDR < 0.1). This reduced the set to 502 genes (**Figure 3A**).

**Figure 3.**
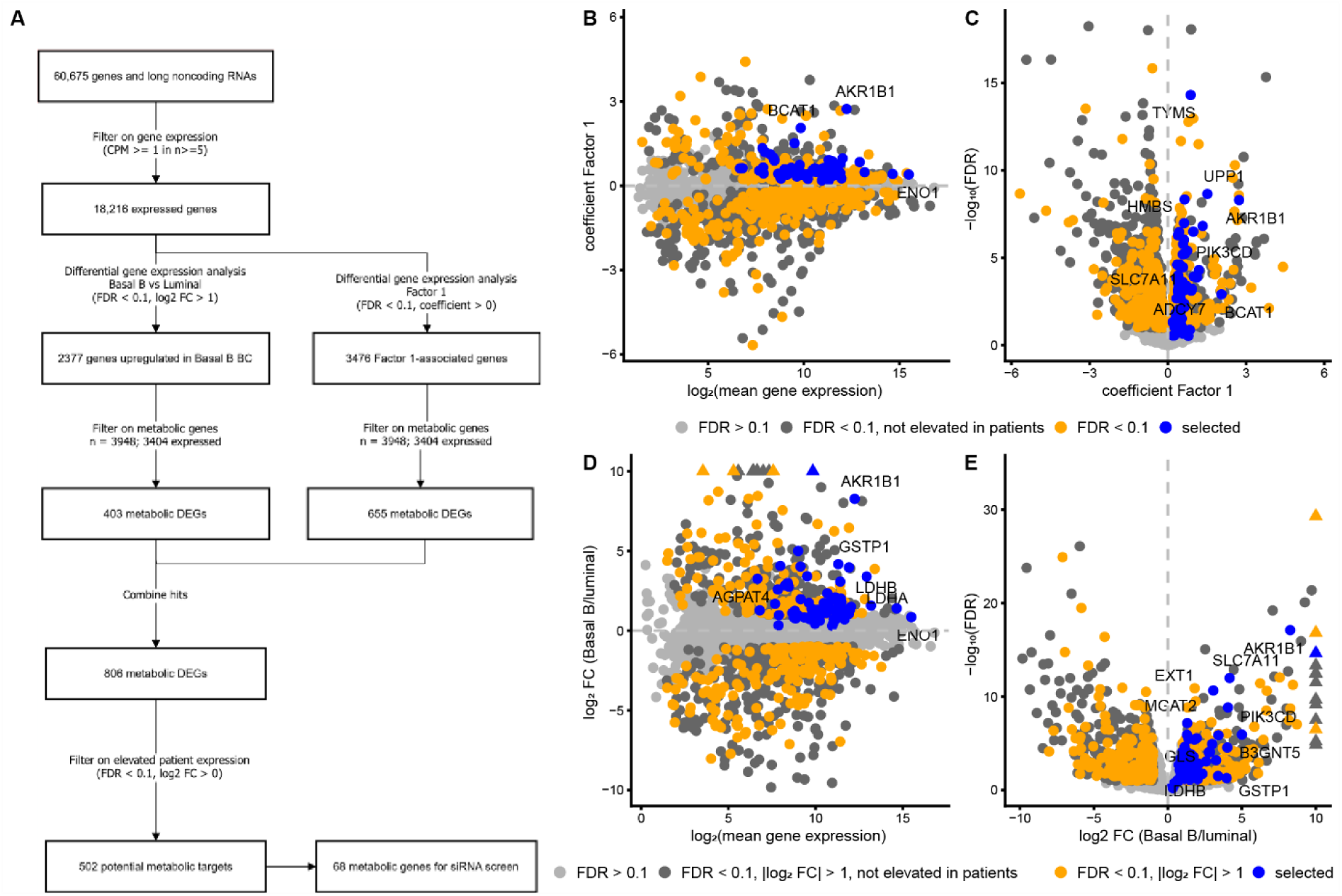
Selection of metabolic genes for a siRNA knockdown screen. (A) Workflow to describe which genes were selected for a knockdown screen. CPM, counts per million; FDR, false discovery rate; BC, breast cancer; DEG, differentially expressed genes; FC, fold change. The four circles with KEGG, Reactome, Recon3D and GO represent the definition of metabolic genes, where genes were selected if they were present in at least two databases. (B-E) MA plots (B, D) and volcano plots (C, E) of genes associated with factor 1 (B, C) or differentially expressed between basal B vs luminal cell lines (D, E).

These selected genes were involved in multiple metabolic sub-processes. This included 68 genes involved in cell cycle regulation (including 20 proteasome components and 32 genes involved in nucleotide metabolism), and 68 genes involved in amino acid metabolism (**Supplementary Figure 3**). Since protein and nucleotide synthesis are required for cell growth, an enhanced expression related to these processes may be linked to the fast-proliferative phenotype of the cell lines. In addition, 10 genes were involved in collagen formation and 19 in glycosaminoglycan metabolism, which are associated with extracellular matrix remodeling and may be linked to migratory and mesenchymal-like properties of the basal B cell lines. Processes such as glycolysis, lipid metabolism, RNA metabolism, and translation (13, 52, 65 and 34 genes, respectively, including 15 ribosomal genes) were also represented. These could be related to both a fast-proliferative or more migratory phenotype. To prioritize novel vulnerabilities, genes with established roles in the cell cycle or RNA synthesis and translation were excluded. We then selected 67 genes, spanning diverse metabolic functions (**Supplementary Figure 4**).

### Knockdown of metabolic genes reduces proliferation and migration

To test whether these metabolic genes influence aggressive phenotypes in breast cancer cell lines, an siRNA knockdown screen was performed in Hs578T cells. Effects on proliferation and cell migration were measured using a SRB assay, live-cell microscopy confluency analysis and a random cell migration assay. Knockdown effects on proliferation were consistent between the two assays (r_s_ = 0.85).

For 34 knockdowns, the proliferation rate was significantly reduced (> 20% decrease; FDR < 0.1) compared to a mock siRNA control in both assays (**Figure 4A-B**). An additional 12 genes reduced proliferation in one assay, and the knockdown of 22 genes did not significantly impact proliferation. Notably, 21 knockdowns reduced proliferation by over 50% in both assays.

**Figure 4.**
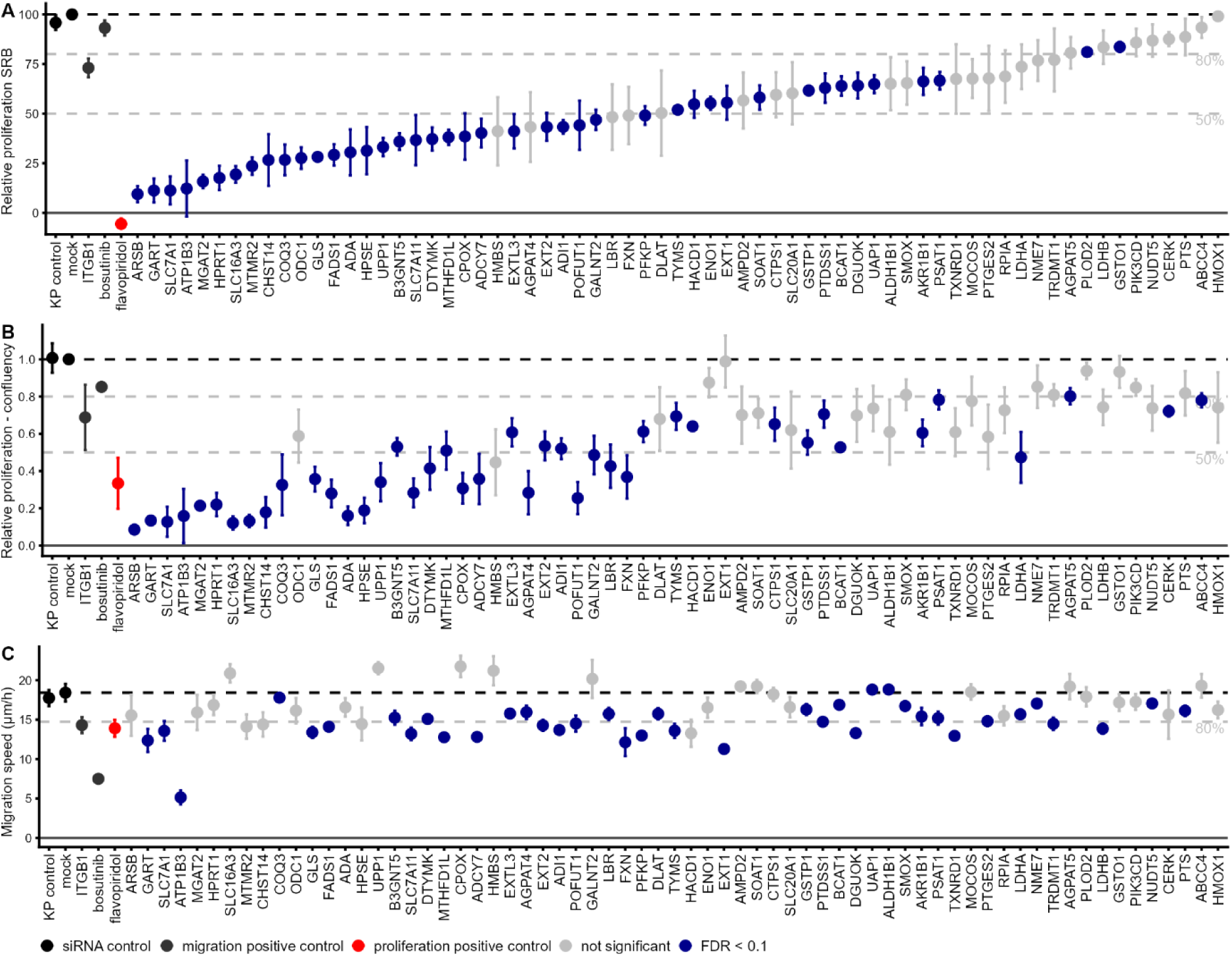
siRNA knockdowns of metabolic targets reduce proliferation and migration in the Hs578T cell line. (A) Relative proliferation of Hs578T after siRNA knockdown compared to the mock siRNA control, measured using an SRB assay. (B) Relative proliferation of Hs578T after siRNA knockdown compared to the mock siRNA control, measured using phase-contrast confluency. The siRNA targets are sorted by their SRB proliferation decrease. (C) Migration speed of Hs578T after siRNA knockdown. The siRNA targets are sorted by their SRB proliferation decrease. All error bars represent the SEM. FDR, false discovery rate. The black dotted line is the value of the mock, and grey dotted lines represent 80% or 50% of this value to highlight a 20-50% decrease.

Twenty knockdowns reduced cell migration by more than 20% (**Figure 4C**). This included genes involved in metabolite transport (ATP1B3, SLC7A1, ADCY7, and SLC7A11), lipid metabolism (FADS1, TXNRD1, and PTDSS1) glycosaminoglycan metabolism (EXT1 and EXT2), nucleotide metabolism (GART, TYMS, DGUOK, TXNRD1), energy metabolism (FXN, and LDHB), amino acid metabolism (GLS, ADI1, TXNRD1) and the genes POFUT1, PFKP, MTHFD1L, and TRDMT1. Eight of these genes also strongly reduced proliferation (>50%, FDR < 0.1), including SLC7A1, MTMR2, GART, ATP1B3, HPSE, POFUT1, FADS1, SLC7A11, GLS, ADCY7, and FXN. In contrast, TXNRD1, LDHB, and TRDMT1 knockdowns reduced migration without significantly affecting proliferation (FDR > 0.1). Of note, the ATP1B3 knockdown reduced the migration speed more than the positive control bosutinib. A detailed inspection of the images showed a low confluency (**Supplementary Figure 5**), suggesting that potential cytotoxicity drove this effect.

Several proliferation- and migration-reducing genes related to metabolic alterations identified earlier. The reduced cell migration after glutaminase (GLS) and the cysteine/glutamate transporter (SCL7A11) knockdowns, both involved in glutamine metabolism, could be related to enhanced glutamine uptake and glutamate release in fast-proliferating, fast-migrating cell lines (**Figure 1F**). The knockdown of the cationic amino acid transporter SLC7A1 could reduce the uptake rate of multiple proliferation-associated amino acids (**Figure 2**), and the knockdown of phosphatidylserine synthase 1 (PTDSS1) could influence intracellular serine and PS levels.

Multiple knockdowns of glycosaminoglycan-related genes substantially reduced proliferation. ARSB reduced proliferation by 90%, and other glycosaminoglycan-associated genes MGAT2, CHST14, and HPSE reduced proliferation as well (79%, 75% and 70%, respectively, FDR < 0.1). EXT1 and EXT2 reduced both proliferation and migration. The EXT2 knockdown also resulted in a lower confluency, but the confluency of EXT1 was similar to the control. When inspected in more detail, the EXT1 knockdown showed a change in phenotype, with an increased cell size and spread morphology. This morphological change superficially resembled the effects seen upon bosutinib treatment (**Supplementary Figure 5**).

Finally, knockdowns of nucleotide metabolism-associated genes GART and HPRT1 significantly reduced proliferation by 90% (FDR < 0.1). Since GART functions in purine biosynthesis and HPRT1 functions in purine salvage, interfering with these two genes likely reduced purine availability, reducing DNA and RNA synthesis crucial for proliferation. This aligns with CRISPR studies showing reduced tumor formation (GART) and tumor growth (HPRT1) in mice (Tran et al., 2024).

In summary, disrupting various metabolic processes reduced proliferation and migration in the aggressive, fast-proliferating Hs578T breast cancer cell line, underscoring the therapeutic potential of targeting metabolic vulnerabilities to prevent breast cancer progression.

## Discussion

In this study, we systematically profiled intracellular metabolites and metabolite exchange rates in breast cancer cell lines. Integrating this data with previously established lipidomic and transcriptomic datasets and with cell line phenotypes resulted in an in-depth analysis of breast cancer cell line metabolism. Most notably, heterogeneity in cancer cell line metabolism was mainly driven by demands for essential amino acids and nucleotides related to proliferation, and by intrinsic differences between luminal and basal cancer cell lines, which overlap with an EMT phenotype, especially in glutamine use. Silencing metabolic genes elevated in a fast-proliferating, basal B, mesenchymal-like cell line reduced both proliferation and migration. In this discussion, we will highlight a subset of these potential drug targets in more detail, ending with limitations of this study and future prospects.

Out of a total of 34 knockdowns reducing proliferation and 20 genes reducing migration, EXT1, EXT2 and metabolite transporters are of interest. Silencing either EXT1 or EXT2 reduced proliferation and migration, and the EXT1 knockdown also altered cell morphology. The glycosyltransferases EXT1 and EXT2 form a heterodimer involved in heparan sulfate synthesis. Targeting both proliferative and migratory phenotypes by interfering with EXT1/EXT2 could be a promising strategy to prevent cancer progression. Although EXT1 was once considered a tumor suppressor (Lind et al., 1998; Ropero et al., 2004), recent studies link it to aggressive cancer (Kong et al., 2021; Manandhar et al., 2017; Solaimuthu et al., 2024) and worse overall survival in triple-negative breast cancer patients (Hossny et al., 2024). Similar to our findings, a knockdown of EXT1 in non-small cell lung carcinoma cell lines reduced proliferation and migration, which the authors attributed to changes in Wnt/β-catenin signaling (Kong et al., 2021). Another study showed that EXT1 was elevated in a doxorubicin-resistant breast cancer cell line MCF7/ADR, and that an EXT1 knockdown reduced the aggressive drug-resistant phenotype and reduced EMT features in these cells (Manandhar et al., 2017). A recent report showed the role of EXT1 in great detail in breast cancer, further highlighting that EXT1 is elevated during EMT, that a knockdown of EXT1 in MDA-MB-231 reduced migration, related to JAK/STAT3 signaling (Solaimuthu et al., 2024). Our findings in Hs578T align with these reports and further strengthening the rationale for targeting EXT1/EXT2.

The knockdown of the metabolite transporter SLC7A1 reduced proliferation and cell migration. Its expression was elevated in aggressive breast cancer cell lines and basal breast cancer patients. Because of its broad role as a cationic metabolite transporter, especially transporting arginine, and because we have shown that aggressive cell lines take up more amino acids including arginine, this transporter could be a promising drug target as well. In NSCLC, SLC7A1 silencing reduced arginine import and reduced cell growth (Banjarnahor et al., 2022; Gai et al., 2024). In epithelial ovarian cancer, SLC7A1 was upregulated in ovarian cancer cells as well as in cancer-associated fibroblasts, and a SLC7A1 knockdown reduced EMT progression and cell migration (You et al., 2024). Recent studies have screened potential inhibitors for SLC7A1, identifying verapamil as an inhibitor of arginine transport in HEK cells (Banjarnahor et al., 2022). In osteosarcoma, inhibiting SLC7A1 reduced proliferation, and cepharanthine was identified as a potential inhibitor (Liao et al., 2025).

More generally, metabolite transporters are attractive because the fast-proliferating aggressive cell lines depend heavily on metabolite uptake. Targeting transporters is showing potential in cancer therapy (Nwosu et al., 2023). In our data, transporter expression patterns varied across subtypes, suggesting that a personalized approach may be required. We have identified three metabolite transporters associated with a slower-proliferating luminal phenotype, SLC7A8, SLC1A4 and SLC38A1. Elevated SLC7A8 has been associated with a favorable prognosis in slow-proliferating ER+ breast cancer (El Ansari et al., 2020), which matches with our findings, whereas SLC1A4 and SLC38A1 were previously associated with cancer proliferation (Hushmandi et al., 2024; K. Wang et al., 2013). Inhibiting any of these three transporters was found to reduce breast cancer growth (van Geldermalsen et al., 2018; K. Wang et al., 2013), leaving their precise role in proliferation unclear.

In contrast, we identified three transporters SLC3A2, SLC7A11, and SLC16A3 as promising drug targets since these transporters were elevated in aggressive breast cancer cell lines. SLC3A2 was previously associated with proliferation and a poor prognosis in breast cancer patients (El Ansari et al., 2018; Furuya et al., 2012; Ichinoe et al., 2021). SLC3A2 forms a complex with SLC7A11 and mediates cysteine/glutamate exchange (Nwosu et al., 2023), which may explain the elevated glutamate secretion observed in fast-proliferative basal B cell lines. Consistent with this, the SLC7A11 knockdown reduced both proliferation and migration. Silencing SLC16A3 reduced proliferation without impacting migration. SLC16A3 is involved in lactate transport and is regulated by the hypoxia-inducible factor 1α (HIF-1α) (Ullah et al., 2006). High expression of SLC16A3 was associated with a poor prognosis in triple-negative breast cancer (Doyen et al., 2014) and has been suggested as a potential therapeutic target, with multiple inhibitors being tested in preclinical studies (Payen et al., 2020; Singh et al., 2023), including combination therapies with immune checkpoint inhibitors (Babl et al., 2023). A phase I clinical trial has been performed targeting SLC3A2 as well, in acute myeloid leukemia using a monoclonal antibody, IGN523 (NCT02040506, (Bixby et al., 2015)). Recently, it was shown that CD8+ T cells can trigger ferroptosis in tumor cells by downregulating SLC3A2/SLC7A11 through interferon γ secretion (W. Wang et al., 2019), which makes immunotherapy a potential alternative strategy to target the function metabolite transporters, next to using small inhibitors or antibodies.

Targeting transporters faces challenges due to adaptive resistance. Based on our data, glutamine import would be a potential target because of the elevated uptake, and others (Jain et al., 2012) have shown this dependency as well in aggressive cancer cells. Glutamine addiction has been studied with much interest in a cancer context (Wise & Thompson, 2010), but metabolic rewiring, leading to drug resistance, is a caveat. Cells can gain resistance against glutamine import inhibition by increasing macropinocytosis (Wahi et al., 2024) or by upregulating glutamine synthase (Tardito et al., 2015). Similarly, cells adapt to a low arginine availability by upregulating arginine synthesis-related genes ASS1 or ASNS (Lukey et al., 2017; Qiu et al., 2015). This means that an effective treatment would require limiting both metabolite availability and synthesis, and this limitation is not unique for amino acid metabolism. For example, reducing purine availability by targeting GART (de novo synthesis) or HPRT1 (purine salvage) reduced cell proliferation in our study, and reductions in tumor growth have been observed by others (Chen et al., 2025; Tran et al., 2024). However, supplementing nucleotides in the diet accelerated tumor growth (Tran et al., 2024), highlighting the complex interplay between nutrient availability and nutrient synthesis. Combining metabolic inhibition with a reduced nutrient availability in the diet shows promising results, for example in neuroblastoma (Cherkaoui et al., 2024), and could have potential in breast cancer as well.

Our study has several limitations. First, all cell lines were cultured in one medium, RPMI, which does not fully reflect tumor interstitial fluid or plasma (Lagziel et al., 2020). As a result, some context-dependent sensitivities may have been missed. Second, we could not fully distinguish between metabolic characteristics caused by the fast-proliferative or by the basal B, mesenchymal-like, phenotype of cancer cell lines, since both phenotypes overlap in our cancer cell line panel. Future studies could induce EMT or a mesenchymal-to-epithelial transition and study resulting changes in the metabolome. A complication will then be that an EMT switch also reduces proliferation. EMT induction should therefore be paired with studies directly inhibiting proliferation, for example by using cell checkpoint inhibitors, in order to reduce the risk of attributing proliferation-associated effects to EMT. Finally, although we have profiled a wide selection of metabolites, we used a targeted metabolomics approach and have not profiled the full metabolome. The cellular metabolome is influenced by a complex network of reactions based on enzyme and nutrient availability, meaning that a network approach may be required to fully understand the impact of metabolic changes. By profiling more metabolites in more specific environmental metabolic contexts, we expect that future studies can obtain a more complete overview of metabolism, leading to promising metabolic targets to treat breast cancer.

## Acknowledgements

This work was supported by the Netherlands Organization for Scientific Research (NWO) Enabling Technology Hotels Program (Project Number 435004026) and the China Scholarship Council (201506220181).

## Supplementary Figures

**Supplementary Figure 1.**
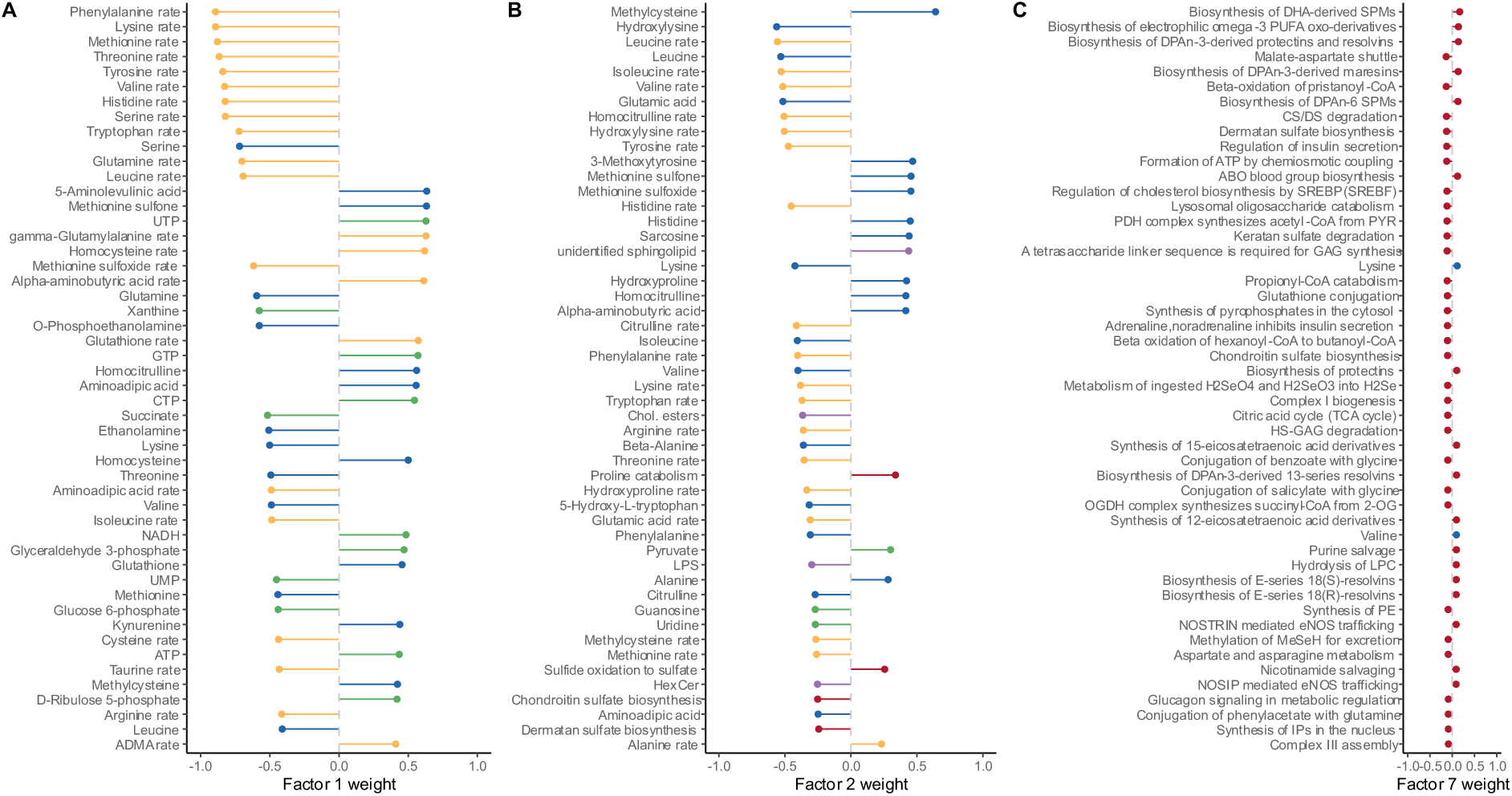
Factor weights of the top 50 features for factor 1, 2, and 7. Blue = amines, yellow = metabolite rates, green = polar metabolites, purple = lipid classes, red = gene expression pathways.

**Supplementary Figure 2.**
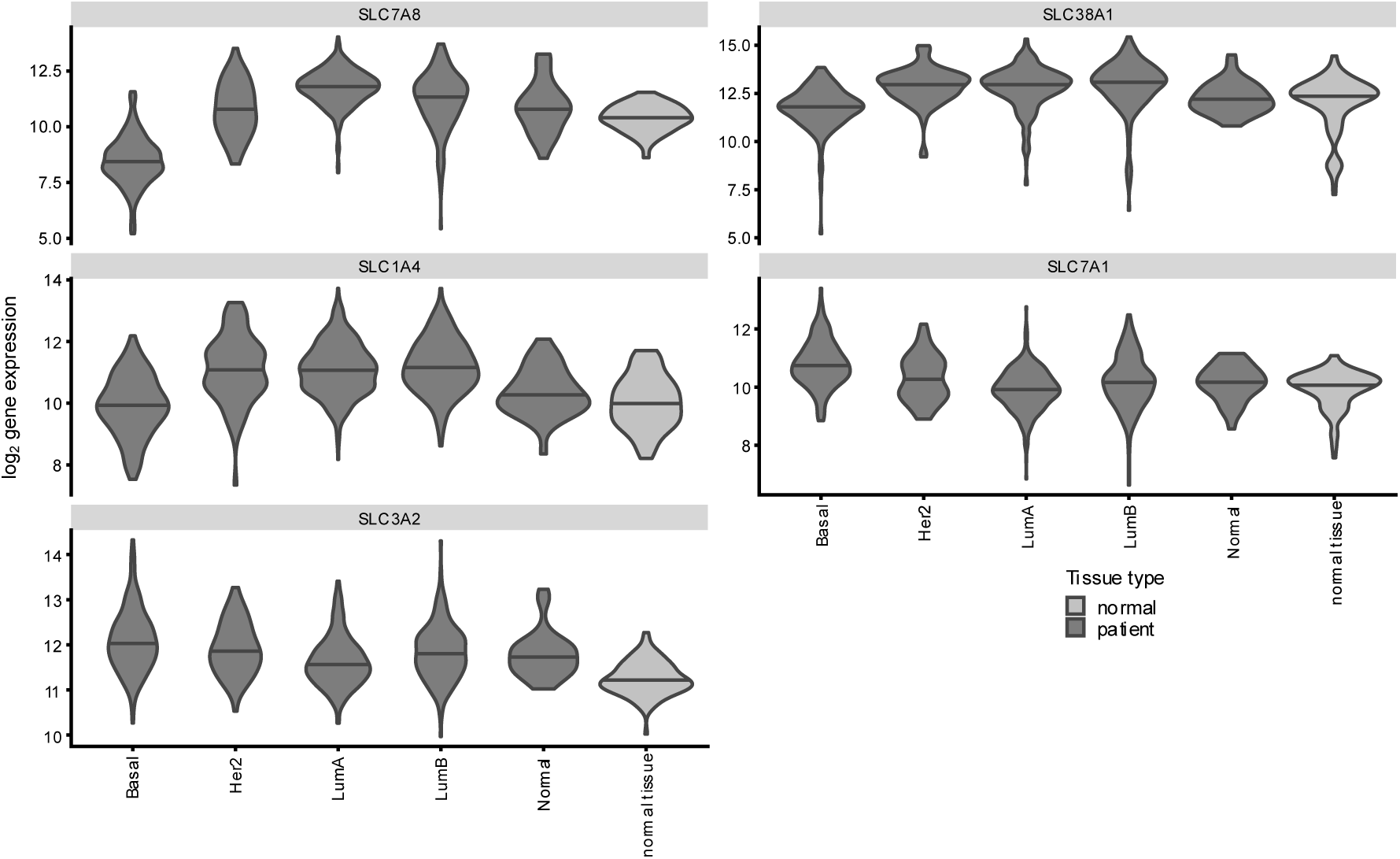
Gene expression profiles of five metabolite transporters in breast cancer patients, split by breast cancer subtype.

**Supplementary Figure 3.**
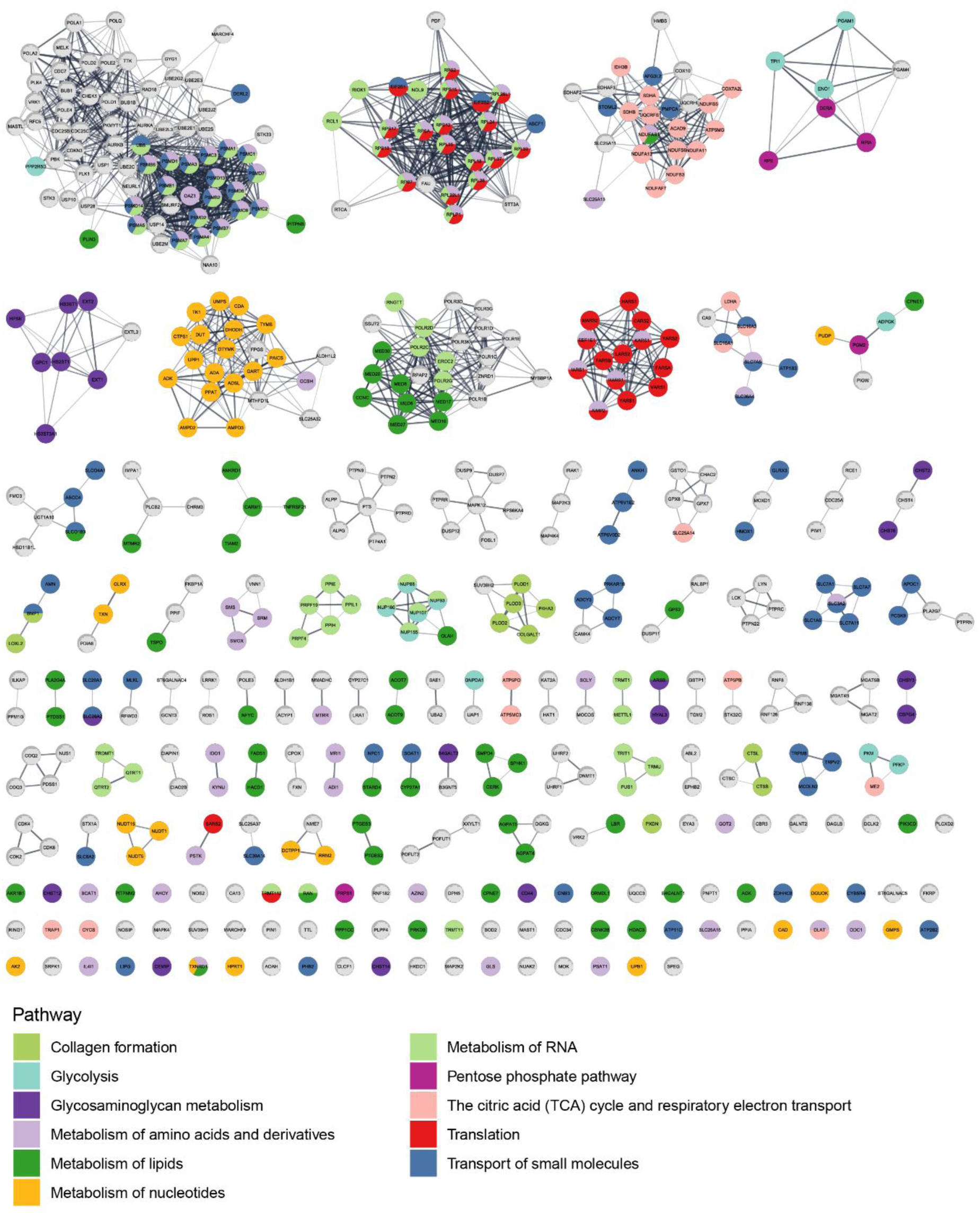
Differentially expressed genes mapped to a STRING-DB protein-protein interaction network, colored by metabolic processes. Grey nodes without a black ring are not part of any of the highlighted pathways. Pathway definitions are from Reactome, or from STRING clusters (Proteasome and cytoplasmic ribosomal proteins)

**Supplementary Figure 4.**
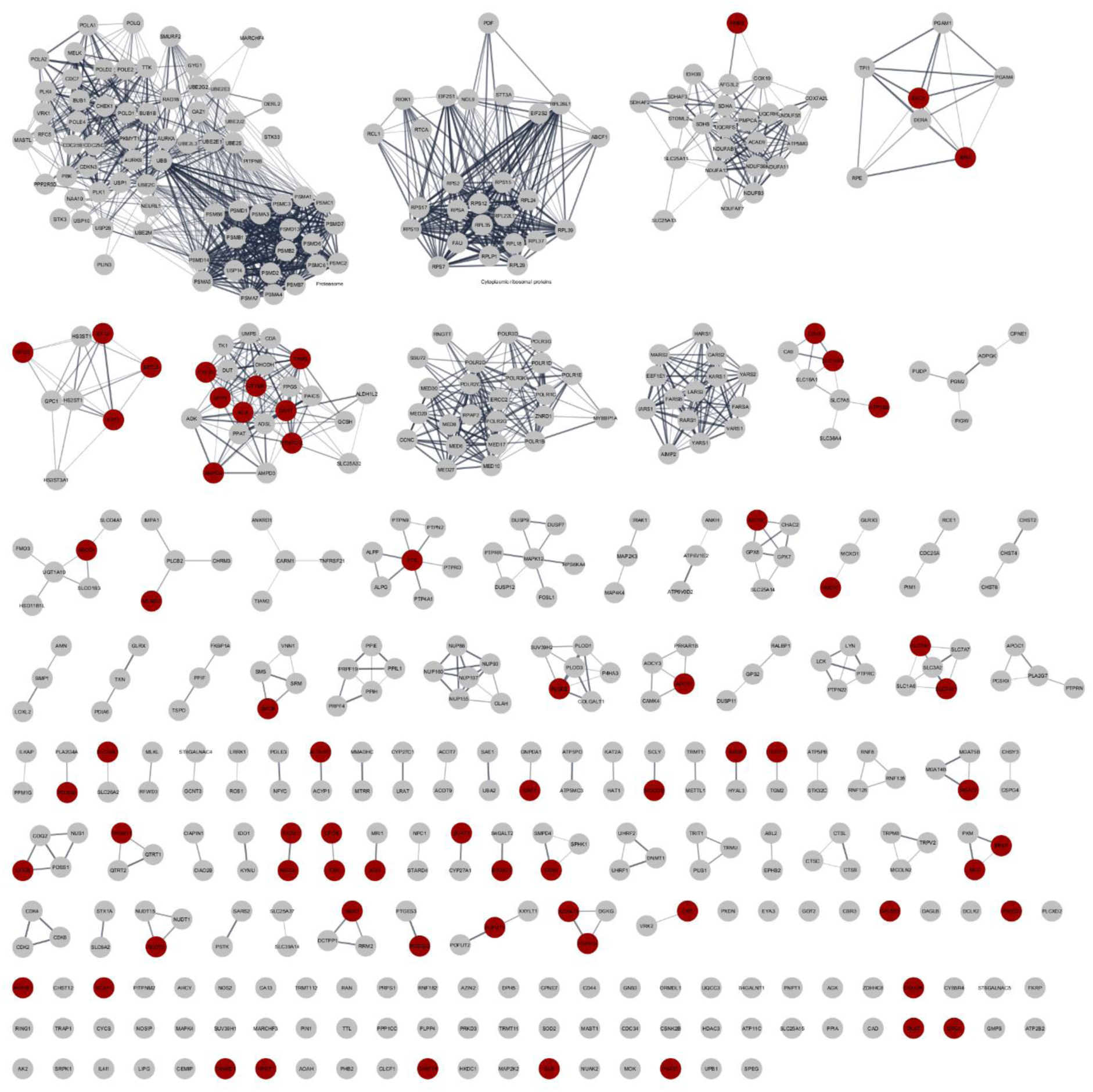
Differentially expressed genes mapped to a STRING-DB protein-protein interaction network. The dark red nodes are selected to target in a siRNA screen.

**Supplementary Figure 5.**
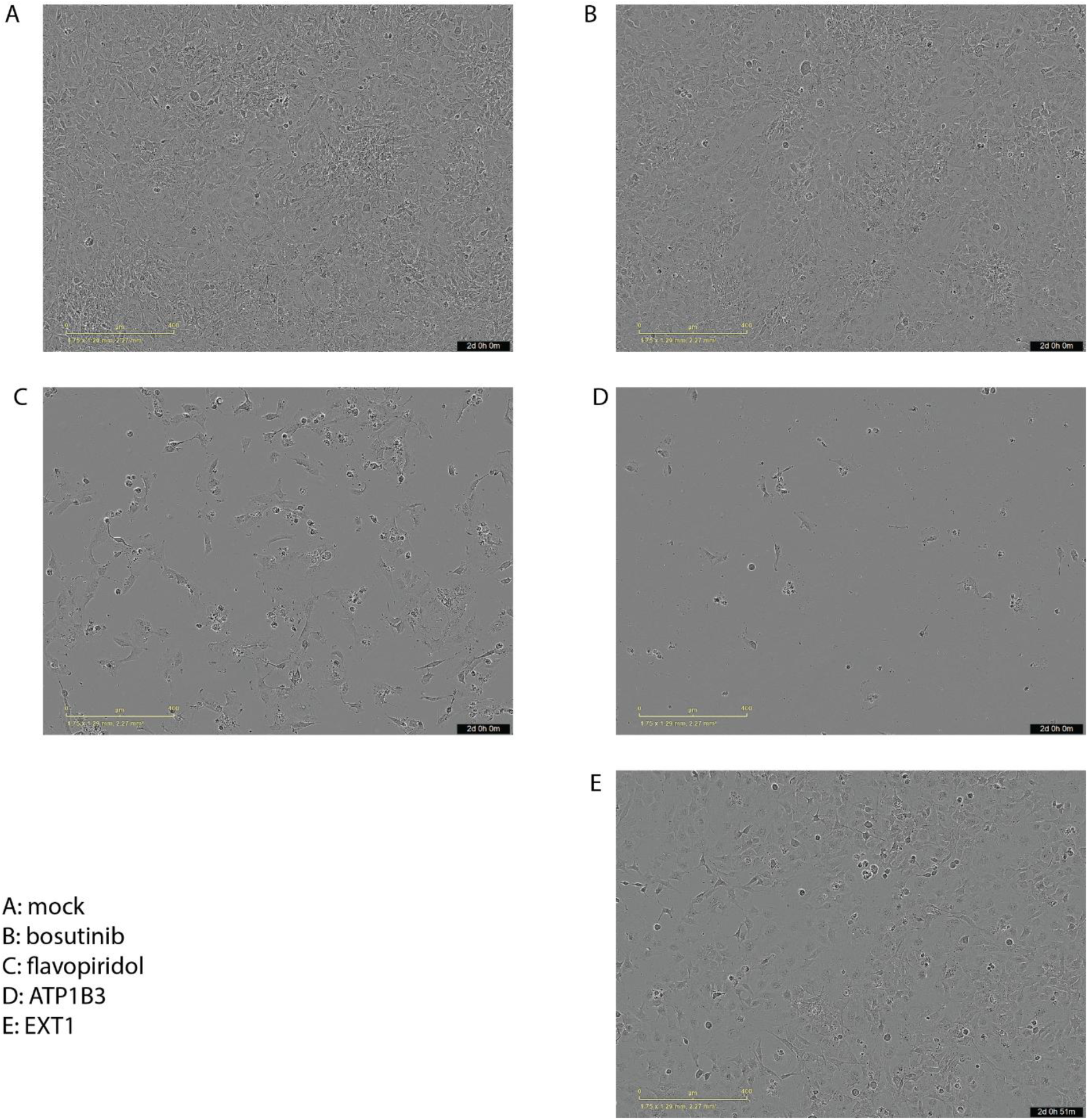
Confluency of cell lines upon siRNA knockdown after two days.

**Supplementary Table 1.** Metabolite abundance in breast cancer cell lines.

*Included as separate file*

## References

Abbott, K. L., Ali, A., Casalena, D., Do, B. T., Ferreira, R., Cheah, J. H., Soule, C. K., Deik, A., Kunchok, T., Schmidt, D. R., Renner, S., Honeder, S. E., Wu, M., Chan, S. H., Tseyang, T., Stoltzfus, A. T., Michel, S. L. J., Greaves, D., Hsu, P. P.,…Vander Heiden, M. G. (2023). Screening in serum-derived medium reveals differential response to compounds targeting metabolism. Cell Chemical Biology, 30(9), 1156–1168.e7. 10.1016/j.chembiol.2023.08.007

Ackermann, T., & Tardito, S. (2019). Cell culture medium formulation and its implications in cancer metabolism. Trends in Cancer, 5(6), 329–332. 10.1016/j.trecan.2019.05.004

Argelaguet, R., Velten, B., Arnol, D., Dietrich, S., Zenz, T., Marioni, J. C., Buettner, F., Huber, W., & Stegle, O. (2018). Multi-Omics Factor Analysis—A framework for unsupervised integration of multi-omics data sets. Molecular Systems Biology, 14(6), 1–13. 10.15252/msb.20178124

Ashburner, M., Ball, C. A., Blake, J. A., Botstein, D., Butler, H., Cherry, J. M., Davis, A. P., Dolinski, K., Dwight, S. S., Eppig, J. T., Harris, M. A., Hill, D. P., Issel-Tarver, L., Kasarskis, A., Lewis, S., Matese, J. C., Richardson, J. E., Ringwald, M., Rubin, G. M., & Sherlock, G. (2000). Gene Ontology: Tool for the unification of biology. Nature Genetics, 25(1), 25–29. 10.1038/75556

Babl, N., Decking, S.-M., Voll, F., Althammer, M., Sala-Hojman, A., Ferretti, R., Korf, C., Schmidl, C., Schmidleithner, L., Nerb, B., Matos, C., Koehl, G. E., Siska, P., Bruss, C., Kellermeier, F., Dettmer, K., Oefner, P. J., Wichland, M., Ugele, I.,…Renner, K. (2023). MCT4 blockade increases the efficacy of immune checkpoint blockade. Journal for ImmunoTherapy of Cancer, 11(10), e007349. 10.1136/jitc-2023-007349

Banjarnahor, S., König, J., & Maas, R. (2022). Screening of commonly prescribed drugs for effects on the CAT1-mediated transport of l-arginine and arginine derivatives. Amino Acids, 54(7), 1101–1108. 10.1007/s00726-022-03156-2

Bergers, G., & Fendt, S.-M. (2021). The metabolism of cancer cells during metastasis. Nature Reviews Cancer, 21(3), 162–180. 10.1038/s41568-020-00320-2

Bixby, D., Wieduwilt, M. J., Akard, L. P., Khoury, H. J., Becker, P. S., Van Der Horst, E. H., Ho, W., & Cortes, J. E. (2015). A Phase I Study of IGN523, a Novel Anti-CD98 Monoclonal Antibody in Patients with Relapsed or Refractory Acute Myeloid Leukemia (AML). Blood, 126(23), 3809–3809. 10.1182/blood.V126.23.3809.3809

Brunk, E., Sahoo, S., Zielinski, D. C., Altunkaya, A., Dräger, A., Mih, N., Gatto, F., Nilsson, A., Preciat Gonzalez, G. A., Aurich, M. K., Prlić, A., Sastry, A., Danielsdottir, A. D., Heinken, A., Noronha, A., Rose, P. W., Burley, S. K., Fleming, R. M. T., Nielsen, J.,…Palsson, B. O. (2018). Recon3D enables a three-dimensional view of gene variation in human metabolism. Nature Biotechnology, 36(3), 272–281. 10.1038/nbt.4072

Cao, M. D., Lamichhane, S., Lundgren, S., Bofin, A., Fjøsne, H., Giskeødegård, G. F., & Bathen, T. F. (2014). Metabolic characterization of triple negative breast cancer. BMC Cancer, 14(1). 10.1186/1471-2407-14-941

Cerami, E., Gao, J., Dogrusoz, U., Gross, B. E., Sumer, S. O., Aksoy, B. A., Jacobsen, A., Byrne, C. J., Heuer, M. L., Larsson, E., Antipin, Y., Reva, B., Goldberg, A. P., Sander, C., & Schultz, N. (2012). The cBio Cancer Genomics Portal: An Open Platform for Exploring Multidimensional Cancer Genomics Data. Cancer Discovery, 2(5), 401–404. 10.1158/2159-8290.CD-12-0095

Chandel, N. S., Vousden, K. H., & DeBerardinis, R. J. (2025). Cancer Metabolism: Historical Landmarks, New Concepts, and Opportunities. Cold Spring Harbor Perspectives in Medicine, 15(12). 10.1101/cshperspect.a041814

Chen, Z., Ding, Y.-H., Zhao, M., Zhang, Y., Sun, M.-Y., Zhang, A.-Q., Qian, X., & Ji, X.-M. (2025). GART promotes the proliferation and migration of human non-small cell lung cancer cell lines A549 and H1299 by targeting PAICS-Akt-β-catenin pathway. Frontiers in Oncology, 15, 1543463. 10.3389/fonc.2025.1543463

Cherkaoui, S., Durot, S., Bradley, J., Critchlow, S., Dubuis, S., Masiero, M. M., Wegmann, R., Snijder, B., Othman, A., Bendtsen, C., & Zamboni, N. (2022). A functional analysis of 180 cancer cell lines reveals conserved intrinsic metabolic programs. Molecular Systems Biology, 18(11), e11033. 10.15252/msb.202211033

Cherkaoui, S., Yang, L., McBride, M., Turn, C. S., Lu, W., Eigenmann, C., Allen, G. E., Panasenko, O. O., Zhang, L., Vu, A., Liu, K., Li, Y., Gandhi, O. H., Surrey, L., Wierer, M., White, E., Rabinowitz, J. D., Hogarty, M. D., & Morscher, R. J. (2024). Reprogramming neuroblastoma by diet-enhanced polyamine depletion. bioRxiv, 2024.01.07.573662. 10.1101/2024.01.07.573662

Dai, X., Cheng, H., Bai, Z., & Li, J. (2017). Breast cancer cell line classification and its relevance with breast tumor subtyping. Journal of Cancer, 8(16), 3131–3141. 10.7150/jca.18457

Demas, D. M., Demo, S., Fallah, Y., Clarke, R., Nephew, K. P., Althouse, S., Sandusky, G., He, W., & Shajahan-Haq, A. N. (2019). Glutamine Metabolism Drives Growth in Advanced Hormone Receptor Positive Breast Cancer. Frontiers in Oncology, 9. 10.3389/fonc.2019.00686

Demicco, M., Liu, X.-Z., Leithner, K., & Fendt, S.-M. (2024). Metabolic heterogeneity in cancer. Nature Metabolism, 6(1), 18–38. 10.1038/s42255-023-00963-z

Dieterle, F., Ross, A., Schlotterbeck, G., & Senn, H. (2006). Probabilistic Quotient Normalization as Robust Method to Account for Dilution of Complex Biological Mixtures. Application in 1H NMR Metabonomics. Analytical Chemistry, 78(13), 4281–4290. 10.1021/AC051632C

Doncheva, N. T., Morris, J. H., Gorodkin, J., & Jensen, L. J. (2019). Cytoscape StringApp: Network Analysis and Visualization of Proteomics Data. Journal of Proteome Research, 18(2), 623–632. 10.1021/acs.jproteome.8b00702

Doyen, J., Trastour, C., Ettore, F., Peyrottes, I., Toussant, N., Gal, J., Ilc, K., Roux, D., Parks, S. K., Ferrero, J. M., & Pouysségur, J. (2014). Expression of the hypoxia-inducible monocarboxylate transporter MCT4 is increased in triple negative breast cancer and correlates independently with clinical outcome. Biochemical and Biophysical Research Communications, 451(1), 54–61. 10.1016/j.bbrc.2014.07.050

El Ansari, R., Alfarsi, L., Craze, M. L., Masisi, B. K., Ellis, I. O., Rakha, E. A., & Green, A. R. (2020). The solute carrier SLC7A8 is a marker of favourable prognosis in ER-positive low proliferative invasive breast cancer. Breast Cancer Research and Treatment, 181(1), 1–12. 10.1007/s10549-020-05586-6

Elia, I., Doglioni, G., & Fendt, S.-M. (2018). Metabolic Hallmarks of Metastasis Formation. Trends in Cell Biology, 28(8), 673–684. 10.1016/j.tcb.2018.04.002

Gai, X., Liu, Y., Lan, X., Chen, L., Yuan, T., Xu, J., Li, Y., Zheng, Y., Yan, Y., Yang, L., Fu, Y., Tang, S., Cao, S., Dai, X., Zhu, H., Geng, M., Ding, J., Pu, C., & Huang, M. (2024). Oncogenic KRAS Induces Arginine Auxotrophy and Confers a Therapeutic Vulnerability to SLC7A1 Inhibition in Non–Small Cell Lung Cancer. Cancer Research, 84(12), 1963–1977. 10.1158/0008-5472.CAN-23-2095

Gu, Z. (2022). Complex heatmap visualization. iMeta, 1(3), e43. 10.1002/imt2.43

Hanahan, D., & Weinberg, R. A. (2011). Hallmarks of cancer: The next generation. Cell, 144(5), 646–674. 10.1016/j.cell.2011.02.013

Heirendt, L., Arreckx, S., Pfau, T., Mendoza, S. N., Richelle, A., Heinken, A., Haraldsdóttir, H. S., Wachowiak, J., Keating, S. M., Vlasov, V., Magnusdóttir, S., Ng, C. Y., Preciat, G., Žagare, A., Chan, S. H. J., Aurich, M. K., Clancy, C. M., Modamio, J., Sauls, J. T.,…Fleming, R. M. T. (2019). Creation and analysis of biochemical constraint-based models using the COBRA Toolbox v.3.0. Nature Protocols, 14(3), 639–702. 10.1038/s41596-018-0098-2

Hoadley, K. A., Yau, C., Hinoue, T., Wolf, D. M., Lazar, A. J., Drill, E., Shen, R., Taylor, A. M., Cherniack, A. D., Thorsson, V., Akbani, R., Bowlby, R., Wong, C. K., Wiznerowicz, M., Sanchez-Vega, F., Robertson, A. G., Schneider, B. G., Lawrence, M. S., Noushmehr, H.,…Laird, P. W. (2018). Cell-of-Origin Patterns Dominate the Molecular Classification of 10,000 Tumors from 33 Types of Cancer. Cell, 173(2), 291–304.e6. 10.1016/j.cell.2018.03.022

Hossny, A., Hassan, H. A. F. M., Fahmy, S. A., Abdelazim, H., Kamel, M. M., Osman, A. H., & Ibrahim, S. A. (2024). EXT1 as an Independent Prognostic Biomarker in Breast Cancer: Its Correlation with Immune Infiltration and Clinicopathological Parameters. Immuno, 5(1), 1. 10.3390/immuno5010001

Hushmandi, K., Einollahi, B., Saadat, S. H., Lee, E. H. C., Farani, M. R., Okina, E., Huh, Y. S., Nabavi, N., Salimimoghadam, S., & Kumar, A. P. (2024). Amino acid transporters within the solute carrier superfamily: Underappreciated proteins and novel opportunities for cancer therapy. Molecular Metabolism, 84, 101952. 10.1016/j.molmet.2024.101952

Jain, M., Nilsson, R., Sharma, S., Madhusudhan, N., Kitami, T., Souza, A. L., Kafri, R., Kirschner, M. W., Clish, C. B., & Mootha, V. K. (2012). Metabolite profiling identifies a key role for glycine in rapid cancer cell proliferation. Science, 336(6084), 1040–1044. 10.1126/science.1218595

Kanehisa, M., Furumichi, M., Sato, Y., Matsuura, Y., & Ishiguro-Watanabe, M. (2024). KEGG: Biological systems database as a model of the real world. Nucleic Acids Research, 53(D1), D672–D677. 10.1093/nar/gkae909

Katzir, R., Polat, I. H., Harel, M., Katz, S., Foguet, C., Selivanov, V. A., Sabatier, P., Cascante, M., Geiger, T., & Ruppin, E. (2019). The landscape of tiered regulation of breast cancer cell metabolism. Scientific Reports, 9(1), 17760. 10.1038/s41598-019-54221-y

Kim, S., Kim, D. H., Jung, W.-H., & Koo, J. S. (2013). Expression of glutamine metabolism-related proteins according to molecular subtype of breast cancer. Endocrine-Related Cancer, 20(3), 339–348. 10.1530/ERC-12-0398

Kleensang, A., Vantangoli, M. M., Odwin-DaCosta, S., Andersen, M. E., Boekelheide, K., Bouhifd, M., Fornace, A. J., Li, H.-H., Livi, C. B., Madnick, S., Maertens, A., Rosenberg, M., Yager, J. D., Zhao, L., & Hartung, T. (2016). Genetic variability in a frozen batch of MCF-7 cells invisible in routine authentication affecting cell function. Scientific Reports, 6(1), 28994. 10.1038/srep28994

Koedoot, E., Fokkelman, M., Rogkoti, V.-M., Smid, M., van de Sandt, I., de Bont, H., Pont, C., Klip, J. E., Wink, S., Timmermans, M. A., Wiemer, E. A. C., Stoilov, P., Foekens, J. A., Le Dévédec, S. E., Martens, J. W. M., & van de Water, B. (2019). Uncovering the signaling landscape controlling breast cancer cell migration identifies novel metastasis driver genes. Nature Communications, 10(1), 1–16. 10.1038/s41467-019-11020-3

Koedoot, E., Wolters, L., Smid, M., Stoilov, P., Burger, G. A., Herpers, B., Yan, K., Price, L. S., Martens, J. W. M., Le Dévédec, S. E., & van de Water, B. (2021). Differential reprogramming of breast cancer subtypes in 3D cultures and implications for sensitivity to targeted therapy. Scientific Reports, 11(1), 7259. 10.1038/s41598-021-86664-7

Kong, W., Chen, Y., Zhao, Z., Zhang, L., Lin, X., Luo, X., Wang, S., Song, Z., Lin, X., Lai, G., & Yu, Z. (2021). EXT1 methylation promotes proliferation and migration and predicts the clinical outcome of non-small cell lung carcinoma via WNT signalling pathway. Journal of Cellular and Molecular Medicine, 25(5), 2609–2620. 10.1111/jcmm.16277

Lagziel, S., Gottlieb, E., & Shlomi, T. (2020). Mind your media. Nature Metabolism, 2(12), 1369–1372. 10.1038/s42255-020-00299-y

Leegwater, H., Zhang, Z., Zhang, X., Hankemeier, T., Harms, A. C., Zweemer, A. J. M., Le Dévédec, S. E., & Kindt, A. (2024). Normalization Strategies for Lipidome Data in Cell Line Panels. Journal of Chemometrics, 39(1), e3636. 10.1002/cem.3636

Leegwater, H., Zhang, Z., Zhang, X., Wang, X., Hankemeier, T., Zweemer, A. J. M., van de Water, B., Danen, E., Hoekstra, M., Harms, A. C., Kindt, A., & Le Dévédec, S. E. (2025). Distinct lipidomic profiles in breast cancer cell lines relate to proliferation and EMT phenotypes. Biochimica et Biophysica Acta (BBA) - Molecular and Cell Biology of Lipids, 1870(7), 159679. 10.1016/j.bbalip.2025.159679

Li, H., Ning, S., Ghandi, M., Kryukov, G. V., Gopal, S., Deik, A., Souza, A., Pierce, K., Keskula, P., Hernandez, D., Ann, J., Shkoza, D., Apfel, V., Zou, Y., Vazquez, F., Barretina, J., Pagliarini, R. A., Galli, G. G., Root, D. E.,…Sellers, W. R. (2019). The landscape of cancer cell line metabolism. Nature Medicine, 25(5), 850–860. 10.1038/s41591-019-0404-8

Liao, Y., Chen, J., Yao, H., Zheng, T., Tu, J., Chen, W., Guo, Z., Zou, Y., Wen, L., & Xie, X. (2025). Single-cell profiling of SLC family transporters: Uncovering the role of SLC7A1 in osteosarcoma. Journal of Translational Medicine, 23(1), 103. 10.1186/s12967-025-06086-1

Lind, T., Tufaro, F., McCormick, C., Lindahl, U., & Lidholt, K. (1998). The Putative Tumor Suppressors EXT1 and EXT2 Are Glycosyltransferases Required for the Biosynthesis of Heparan Sulfate. Journal of Biological Chemistry, 273(41), 26265–26268. 10.1074/jbc.273.41.26265

Liu, J., Lichtenberg, T., Hoadley, K. A., Poisson, L. M., Lazar, A. J., Cherniack, A. D., Kovatich, A. J., Benz, C. C., Levine, D. A., Lee, A. V., Omberg, L., Wolf, D. M., Shriver, C. D., Thorsson, V., Hu, H., Caesar-Johnson, S. J., Demchok, J. A., Felau, I., Kasapi, M.,…Mariamidze, A. (2018). An integrated TCGA pan-cancer clinical data resource to drive high-quality survival outcome analytics. Cell, 173(2), 400–416.e11. 10.1016/j.cell.2018.02.052

Lopez, M. J., & Mohiuddin, S. S. (2025). Biochemistry, Essential Amino Acids. In StatPearls. StatPearls Publishing. http://www.ncbi.nlm.nih.gov/books/NBK557845/

Lukey, M. J., Katt, W. P., & Cerione, R. A. (2017). Targeting amino acid metabolism for cancer therapy. Drug Discovery Today, 22(5), 796–804. 10.1016/j.drudis.2016.12.003

Manandhar, S., Kim, C.-G., Lee, S.-H., Kang, S. H., Basnet, N., & Lee, Y. M. (2017). Exostosin 1 regulates cancer cell stemness in doxorubicin-resistant breast cancer cells. Oncotarget, 8(41), 70521–70537. 10.18632/oncotarget.19737

Milacic, M., Beavers, D., Conley, P., Gong, C., Gillespie, M., Griss, J., Haw, R., Jassal, B., Matthews, L., May, B., Petryszak, R., Ragueneau, E., Rothfels, K., Sevilla, C., Shamovsky, V., Stephan, R., Tiwari, K., Varusai, T., Weiser, J.,…D’Eustachio, P. (2023). The Reactome Pathway Knowledgebase 2024. Nucleic Acids Research, 52(D1), D672–D678. 10.1093/nar/gkad1025

Morris, J. H., Apeltsin, L., Newman, A. M., Baumbach, J., Wittkop, T., Su, G., Bader, G. D., & Ferrin, T. E. (2011). clusterMaker: A multi-algorithm clustering plugin for Cytoscape. BMC Bioinformatics, 12(1), 436. 10.1186/1471-2105-12-436

Noga, M. J., Dane, A., Shi, S., Attali, A., van Aken, H., Suidgeest, E., Tuinstra, T., Muilwijk, B., Coulier, L., Luider, T., Reijmers, T. H., Vreeken, R. J., & Hankemeier, T. (2011). Metabolomics of cerebrospinal fluid reveals changes in the central nervous system metabolism in a rat model of multiple sclerosis. Metabolomics, 8(2), 253–263. 10.1007/s11306-011-0306-3

Nwosu, Z. C., Song, M. G., di Magliano, M. P., Lyssiotis, C. A., & Kim, S. E. (2023). Nutrient transporters: Connecting cancer metabolism to therapeutic opportunities. Oncogene, 42(10), 711–724. 10.1038/s41388-023-02593-x

Osborne, C. K., Hobbs, K., & Trent, J. M. (1987). Biological differences among MCF-7 human breast cancer cell lines from different laboratories. Breast Cancer Research and Treatment, 9(2), 111–121. 10.1007/BF01807363

Pavlova, N. N., & Thompson, C. B. (2016). The Emerging Hallmarks of Cancer Metabolism. Cell Metabolism, 23(1), 27–47. 10.1016/j.cmet.2015.12.006

Pavlova, N. N., Zhu, J., & Thompson, C. B. (2022). The hallmarks of cancer metabolism: Still emerging. Cell Metabolism, 34(3), 355–377. 10.1016/j.cmet.2022.01.007

Payen, V. L., Mina, E., Van Hée, V. F., Porporato, P. E., & Sonveaux, P. (2020). Monocarboxylate transporters in cancer. Molecular Metabolism, 33, 48–66. 10.1016/j.molmet.2019.07.006

Porcari, A. M., Zhang, J., Garza, K. Y., Rodrigues-Peres, R. M., Lin, J. Q., Young, J. H., Tibshirani, R., Nagi, C., Paiva, G. R., Carter, S. A., Sarian, L. O., Eberlin, M. N., & Eberlin, L. S. (2018). Multicenter study using desorption-electrospray-ionization-mass-spectrometry imaging for breast-cancer diagnosis. Analytical Chemistry, 90(19), 11324–11332. 10.1021/acs.analchem.8b01961

Preciat, G., Wegrzyn, A. B., Moreno, E. L., Willacey, C. C. W., Modamio, J., Monteiro, F. L., El Assal, D., Schurink, A., Oliveira, M. A. P., Zhang, Z., Cousins, B., Haraldsdóttir, H. S., Hachi, S., Zach, S., Leparc, G., Lee, Y. T., Hengerer, B., Vempala, S., Saunders, M. A.,…Fleming, R. M. T. (2021). Mechanistic model-driven exometabolomic characterisation of human dopaminergic neuronal metabolism. Cold Spring Harbor Laboratory. 10.1101/2021.06.30.450562

Qiu, F., Huang, J., & Sui, M. (2015). Targeting arginine metabolism pathway to treat arginine-dependent cancers. Cancer Letters, 364(1), 1–7. 10.1016/j.canlet.2015.04.020

R Core Team. (2024). R: a language and environment for statistical computing [Manual]. R Foundation for Statistical Computing. https://www.R-project.org/

Rogkoti, V. M., Angelopoulos, N., Bont, H. de, Smid, M., Klop, M., Di, Z., Meerman, J. H. N., Foekens, J. A., Martens, J. W. M., Price, L. S., Wessels, L. F. A., Water, B. van de, & Dévédec, S. E. L. (n.d.). An integrated systems microscopy and transcriptomics analysis identifies gene signatures of breast cancer cell migratory and invasive behavior. Submitted.

Ropero, S., Setien, F., Espada, J., Fraga, M. F., Herranz, M., Asp, J., Benassi, M. S., Franchi, A., Patiño, A., Ward, L. S., Bovee, J., Cigudosa, J. C., Wim, W., & Esteller, M. (2004). Epigenetic loss of the familial tumor-suppressor gene exostosin-1 (EXT1) disrupts heparan sulfate synthesis in cancer cells. Human Molecular Genetics, 13(22), 2753–2765. 10.1093/hmg/ddh298

Rossi, M., Altea-Manzano, P., Demicco, M., Doglioni, G., Bornes, L., Fukano, M., Vandekeere, A., Cuadros, A. M., Fernández-García, J., Riera-Domingo, C., Jauset, C., Planque, M., Alkan, H. F., Nittner, D., Zuo, D., Broadfield, L. A., Parik, S., Pane, A. A., Rizzollo, F.,…Fendt, S.-M. (2022). PHGDH heterogeneity potentiates cancer cell dissemination and metastasis. Nature, 605(7911), 747–753. 10.1038/s41586-022-04758-2

Shaul, Y. D., Freinkman, E., Comb, W. C., Cantor, J. R., Tam, W. L., Thiru, P., Kim, D., Kanarek, N., Pacold, M. E., Chen, W. W., Bierie, B., Possemato, R., Reinhardt, F., Weinberg, R. A., Yaffe, M. B., & Sabatini, D. M. (2014). Dihydropyrimidine accumulation is required for the epithelial-mesenchymal transition. Cell, 158(5), 1094–1109. 10.1016/j.cell.2014.07.032

Singh, M., Afonso, J., Sharma, D., Gupta, R., Kumar, V., Rani, R., Baltazar, F., & Kumar, V. (2023). Targeting monocarboxylate transporters (MCTs) in cancer: How close are we to the clinics? Seminars in Cancer Biology, 90, 1–14. 10.1016/j.semcancer.2023.01.007

Solaimuthu, B., Khatib, A., Tanna, M., Karmi, A., Hayashi, A., Abu Rmaileh, A., Lichtenstein, M., Takoe, S., Jolly, M. K., & Shaul, Y. D. (2024). The exostosin glycosyltransferase 1/STAT3 axis is a driver of breast cancer aggressiveness. Proceedings of the National Academy of Sciences, 121(3), e2316733121. 10.1073/pnas.2316733121

Szklarczyk, D., Kirsch, R., Koutrouli, M., Nastou, K., Mehryary, F., Hachilif, R., Gable, A. L., Fang, T., Doncheva, N. T., Pyysalo, S., Bork, P., Jensen, L. J., & von Mering, C. (2023). The STRING database in 2023: Protein–protein association networks and functional enrichment analyses for any sequenced genome of interest. Nucleic Acids Research, 51(D1), D638–D646. 10.1093/nar/gkac1000

Tan, J., & Le, A. (2021). The Heterogeneity of Breast Cancer Metabolism. In A. Le (Ed.), The Heterogeneity of Cancer Metabolism (pp. 89–101). Springer International Publishing. 10.1007/978-3-030-65768-0_6

Tardito, S., Oudin, A., Ahmed, S. U., Fack, F., Keunen, O., Zheng, L., Miletic, H., Sakariassen, P. Ø., Weinstock, A., Wagner, A., Lindsay, S. L., Hock, A. K., Barnett, S. C., Ruppin, E., Mørkve, S. H., Lund-Johansen, M., Chalmers, A. J., Bjerkvig, R., Niclou, S. P., & Gottlieb, E. (2015). Glutamine synthetase activity fuels nucleotide biosynthesis and supports growth of glutamine-restricted glioblastoma. Nature Cell Biology, 17(12), 1556–1568. 10.1038/ncb3272

The Gene Ontology Consortium, Aleksander, S. A., Balhoff, J., Carbon, S., Cherry, J. M., Drabkin, H. J., Ebert, D., Feuermann, M., Gaudet, P., Harris, N. L., Hill, D. P., Lee, R., Mi, H., Moxon, S., Mungall, C. J., Muruganugan, A., Mushayahama, T., Sternberg, P. W., Thomas, P. D.,…Westerfield, M. (2023). The Gene Ontology knowledgebase in 2023. GENETICS, 224(1), iyad031. 10.1093/genetics/iyad031

Tran, D. H., Kim, D., Kesavan, R., Brown, H., Dey, T., Soflaee, M. H., Vu, H. S., Tasdogan, A., Guo, J., Bezwada, D., Al Saad, H., Cai, F., Solmonson, A., Rion, H., Chabatya, R., Merchant, S., Manales, N. J., Tcheuyap, V. T., Mulkey, M.,…Hoxhaj, G. (2024). De novo and salvage purine synthesis pathways across tissues and tumors. Cell, 187(14), 3602–3618.e20. 10.1016/j.cell.2024.05.011

Ullah, M. S., Davies, A. J., & Halestrap, A. P. (2006). The Plasma Membrane Lactate Transporter MCT4, but Not MCT1, Is Up-regulated by Hypoxia through a HIF-1α-dependent Mechanism. Journal of Biological Chemistry, 281(14), 9030–9037. 10.1074/jbc.M511397200

van der Noord, V. E., van der Stel, W., Louwerens, G., Verhoeven, D., Kuiken, H. J., Lieftink, C., Grandits, M., Ecker, G. F., Beijersbergen, R. L., Bouwman, P., Le Dévédec, S. E., & van de Water, B. (2023). Systematic screening identifies ABCG2 as critical factor underlying synergy of kinase inhibitors with transcriptional CDK inhibitors. Breast Cancer Research, 25, 51. 10.1186/s13058-023-01648-x

Van Der Peet, M., Maas, P., Wegrzyn, A., Lamont, L., Fleming, R., Harms, A., Hankemeier, T., & Kindt, A. (2025). mzQuality: A tool for quality monitoring and reporting of targeted mass spectrometry measurements. 10.1101/2025.01.22.633547

van Geldermalsen, M., Quek, L.-E., Turner, N., Freidman, N., Pang, A., Guan, Y. F., Krycer, J. R., Ryan, R., Wang, Q., & Holst, J. (2018). Benzylserine inhibits breast cancer cell growth by disrupting intracellular amino acid homeostasis and triggering amino acid response pathways. BMC Cancer, 18(1), 689. 10.1186/s12885-018-4599-8

Vichai, V., & Kirtikara, K. (2006). Sulforhodamine B colorimetric assay for cytotoxicity screening. Nature Protocols, 1(3), 1112–1116. 10.1038/nprot.2006.179

Wahi, K., Freidman, N., Wang, Q., Devadason, M., Quek, L.-E., Pang, A., Lloyd, L., Larance, M., Zanini, F., Harvey, K., O’Toole, S., Guan, Y. F., & Holst, J. (2024). Macropinocytosis mediates resistance to loss of glutamine transport in triple-negative breast cancer. The EMBO Journal, 43(23), 5857–5882. 10.1038/s44318-024-00271-6

Wang, K., Cao, F., Fang, W., Hu, Y., Chen, Y., Ding, H., & Yu, G. (2013). Activation of SNAT1/SLC38A1 in human breast cancer: Correlation with p-Akt overexpression. BMC Cancer, 13(1), 343. 10.1186/1471-2407-13-343

Wang, W., Green, M., Choi, J. E., Gijón, M., Kennedy, P. D., Johnson, J. K., Liao, P., Lang, X., Kryczek, I., Sell, A., Xia, H., Zhou, J., Li, G., Li, J., Li, W., Wei, S., Vatan, L., Zhang, H., Szeliga, W.,…Zou, W. (2019). CD8+ T cells regulate tumour ferroptosis during cancer immunotherapy. Nature, 569(7755), 270–274. 10.1038/s41586-019-1170-y

Wickham, H. (2016). ggplot2: Elegant Graphics for Data Analysis. Springer-Verlag New York. https://ggplot2.tidyverse.org

Wilke, C. O. (2020). cowplot: Streamlined Plot Theme and Plot Annotations for “ggplot2” [Computer software]. https://CRAN.R-project.org/package=cowplot

Wise, D. R., DeBerardinis, R. J., Mancuso, A., Sayed, N., Zhang, X.-Y., Pfeiffer, H. K., Nissim, I., Daikhin, E., Yudkoff, M., McMahon, S. B., & Thompson, C. B. (2008). Myc regulates a transcriptional program that stimulates mitochondrial glutaminolysis and leads to glutamine addiction. Proceedings of the National Academy of Sciences, 105(48), 18782–18787. 10.1073/pnas.0810199105

Wise, D. R., & Thompson, C. B. (2010). Glutamine addiction: A new therapeutic target in cancer. Trends in Biochemical Sciences, 35(8), 427–433. 10.1016/j.tibs.2010.05.003

You, S., Han, X., Xu, Y., Sui, L., Song, K., & Yao, Q. (2024). High expression of SLC7A1 in high-grade serous ovarian cancer promotes tumor progression and is involved in MAPK/ERK pathway and EMT. Cancer Medicine, 13(10), e7217. 10.1002/cam4.7217

Zaal, E. A., & Berkers, C. R. (2018). The influence of metabolism on drug response in cancer. Frontiers in Oncology, 8, 500. 10.3389/fonc.2018.00500

